# Temporal coupling of field potentials and action potentials in the neocortex

**DOI:** 10.1101/214650

**Authors:** Brendon O. Watson, Mingxin Ding, György Buzsáki

## Abstract

The local field potential (LFP) is an aggregate measure of group neuronal activity and is often correlated with the action potentials of single neurons. In recent years investigators have found that action potential firing rates increase during elevations in power high-frequency band oscillations (50-200 Hz range). However action potentials also contribute to the LFP signal itself, making the spike–LFP relationship complex. Here we examine the relationship between spike rates and LFPs in varying frequency bands in rat neocortical recordings. We find that 50-180Hz oscillations correlate most consistently with high firing rates, but that other LFPs bands also carry information relating to spiking, including in some cases anti-correlations. Relatedly, we find that spiking itself and electromyographic activity contribute to LFP power in these bands. The relationship between spike rates and LFP power varies between brain states and between individual cells. Finally, we create an improved oscillation-based predictor of action potential activity by specifically utilizing information from across the entire recorded frequency spectrum of LFP. The findings illustrate both caveats and improvements to be taken into account in attempts to infer spiking activity from LFP.

## Introduction

Understanding how the local field potential (LFP) relates to spiking activity can yield data of interest both for clinical and scientific purposes (Jacobs & Kahana, 2009; Ray & Maunsell, 2011; Weiss *et al.*, 2013). The LFP reflects the coordinated transmembrane currents summed across nearby neurons, including spikes (Buzsáki *et al.*, 2012; Einevoll *et al.*, 2013) and also has been shown in many brain regions to coordinate the timing of action potential generation (Buzsáki, 2010). Volume-conducted potentials, such as electromyographic signals (EMG), can also part of the compound LFP.

In the cortex in particular large-scale rhythms are less prominent during waking behavior than in other regions such as the hippocampus, with power in the “gamma” band of frequencies (30-200Hz) some of the most prominent during the waking state (Bragin *et al.*, 1995; Watson *et al.*, 2016). As a result, much attention has been paid in recent years to the role of gamma-band (30-100 Hz) activity (Buzsáki & Wang, 2012; Lisman & Jensen, 2013), with several groups extending their analyses to 'high gamma' (or ‘epsilon’ band (Buzsáki & Wang, 2012)) signals of varying definitions, ranging from 60-500 Hz (Canolty *et al.*, 2006; Crone *et al.*, 2006; Colgin *et al.*, 2009; Gaona *et al.*, 2011). In the hippocampus, wavelet analysis identified three distinct gamma bands with each sub-band associated with varying degrees of spike modulation of both pyramidal cells and interneurons (Tort *et al.*, 2010; Belluscio *et al.*, 2012; Schomburg *et al.*, 2014; Fernandez-Ruiz *et al.*, 2017; Lasztóczi & Klausberger, 2017). The precise boundaries between distinct gamma sub-bands and indeed other LFP frequency bands remain less well defined in the neocortex and would benefit from increased exploration.

Indeed, the multitude of forms and diverse characteristic frequencies seen across brain region, species, network state and even cycle-to-cycle variance in excitatory-inhibitory balance (Atallah & Scanziani, 2009; Buzsáki & Wang, 2012) renders the term ‘gamma band activity’ difficult to precisely define. Gamma-band LFP can reflect both local processes and projected patterns from distant regions (Schomburg *et al.*, 2014; Bastos *et al.*, 2015; Fernandez-Ruiz *et al.*, 2017). Sometimes, increased gamma band power is used synonymously with ‘desynchronization’ of the LFP, implying an uncoordinated discharge of pyramidal cells (Renart *et al.*, 2010; Vyazovskiy & Harris, 2013).

Nevertheless, elevated asynchronous multi-unit activity (MUA) activity can also increase spectral power, causing a ‘broadband’ shift in the frequency spectrum (Manning *et al.*, 2009; Ray & Maunsell, 2010, 2011; Belluscio *et al.*, 2012; Lachaux *et al.*, 2012; Scheffer-Teixeira *et al.*, 2013; Miller *et al.*, 2014, 2016; Johnson & Knight, 2015). Other times, gamma band activity is assumed to reflect a true oscillation, supporting temporal coordination and synchronization of principal cells within and across networks (Buzsáki *et al.*, 1983; Gray & Singer, 1989; Bragin *et al.*, 1995; Buzsáki & Chrobak, 1995; Engel *et al.*, 2001; Varela *et al.*, 2001; Fries, 2005). In general, the periodicity of gamma oscillations is thought to reflect a relatively balanced coordination between excitation and inhibition in local cortical circuits (Buzsáki & Wang, 2012). Yet the details of excitation-inhibition coupling are not completely explored and have not been measured at the scale of large populations of neurons.

Over the past decade, many papers have been published taking the pragmatic approach of using localized high gamma as a compound measure for local brain activity, especially in intracranial recordings in patients (Lachaux *et al.*, 2012; Voytek & Knight, 2015). This work is based on well-documented correlates between cortical unit activity and broadband high gamma power in ranges spanning from 40-130Hz (Nir *et al.*, 2007, 2008) to full broad band spectral activity (Miller *et al.*, 2014). Gamma activity is modulated by task and other cognitive aspects (Nir *et al.*, 2007, 2008) and also correlates with BOLD signal (Logothetis *et al.*, 2001), though it remains to be identified whether the energy costs underlying the BOLD is due to inhibitory/excitatory balance driving the oscillations or the spike themselves (Mukamel, 2005; Magri *et al.*, 2012).

Despite progress, the relationship between spiking of neocortical neurons and cortical oscillations remains incompletely defined. It is tempting to use the hippocampal theta oscillation as an instructive example to inform our thinking on any and all oscillations: the theta oscillation in hippocampus is largely externally driven and imposed on local hippocampal neurons, is high amplitude, regular, narrow-band, and is known to coordinate not only spike timing relative to the oscillation but also to each other (Buzsáki, 2010; Colgin, 2016). Yet in the case of cortical gamma band LFP, oscillations are often much lower amplitude (even when corrected for 1/frequency-related power losses observed across the LFP spectrum) less regular, transient, often wider band and it is not clear they have the elaborate effects upon population spiking that hippocampal theta does (Buzsáki & Moser, 2013). Indeed, not all oscillations are equal and in the case of cortical gamma LFP, power may result from occasional genuine narrow band oscillations or may actually be due to simultaneous mixed oscillatory states, EMG or spiking activity itself (Jarvis & Mitra, 2001; Crone *et al.*, 2006; Whittingstall & Logothetis, 2009; Ray & Maunsell, 2011; Belluscio *et al.*, 2012). This is especially important for higher frequencies because the extracellular units reflect not only the short spikes but also the spike-afterdepolarization and - hyperpolarization components of both pyramidal neurons and interneurons (King *et al.*, 1999) can also contribute to various sub-gamma bands (Penttonen *et al.*, 1998). Additionally, both periodic and wide-band components of gamma oscillations can be phase-power modulated by slower rhythms (Buzsáki *et al.*, 1983; Soltesz & Deschênes, 1993; Bragin *et al.*, 1995; Canolty *et al.*, 2006). Although spike contamination of oscillatory power can be a theoretical nuisance, studying the temporal features of such high-frequency events may provide clues about oscillatory events that modulate them, even in situations when invasive unit recordings are not an option. Furthermore, detailed study of spike-LFP coupling with high neuron counts are difficult in patients and may be complimented by animal models wherein larger numbers of neurons can be measured with high-density probes to allow for more detailed quantification of relationships between neurons and the surrounding local field potential.

In the present experiments, we use data recorded from the cortex of rats to examine more precisely the frequency bands, temporal precision and single-cell correlates of spike-LFP coupling across wake, REM (Rapid Eye Movement) sleep and nonREM sleep. We use a data-driven approach to broadly examine the correlates between various frequency bands and diverse neuronal types across many states. We also examine the effects of EMG and spikes themselves on cortical high frequency LFP bands. While we confirm that the power of 50-180Hz oscillations couple positively with population-wide spike rates during waking states we find that this effect is modulated across neurons, states, muscular movement state and even the ratio of excitatory to inhibitory neuron activity. We also find that population spiking activity within the wake state is best predicted using an approach integrating information from across the power spectrum rather than from a single band.

## METHODS

All data and procedures were reported previously (Watson *et al.*, 2016). All protocols were approved by the Institutional Animal Care and Use Committee of New York University and Weill Cornell Medical College.

Briefly, subjects were male Long-Evans rats aged 3-7 months and were recorded during free wake and sleep behavior in their home cages at least 7 days after chronic surgical implantation of silicon probes. Data was acquired at 20 kHz and was separately low pass filtered to 1250Hz for local field potential (LFP) signal and high pass filtered at 800 Hz and thresholded at 3.5 standard deviations below the mean to find spiking events using the NDManager Suite (Hazan *et al.*, 2006). Threshold crossings which were then grouped into clusters using KlustaKwik (Rossant *et al.*, 2016) and manually screened using klusters (Hazan *et al.*, 2006).

The dataset includes 11 animals and 27 recording sessions, 995 putative excitatory (pE) units and 126 putative inhibitory (pI) units based on classification using spike waveforms shape (Stark *et al.*, 2013; Watson *et al.*, 2016). In brief, pE units had wider waveforms and pI units had narrower waveforms as measured by a combination of trough-to-peak times and inverse of maximal wavelet frequency. This separation method was generated based on optogenetically-tagged parvalbumin positive pI units and CamKII-positive pE units (Stark et al, 2013). Further justification comes from work showing that pE units tended to be followed by increases in spiking by surrounding units, at 1-5ms time lag while pI units are followed by decreases in spiking by surrounding units at the same time lag (Watson et al, 2016; Supplemental Figure 2A).

EMG-related signal was extracted from the LFP signal using correlations between distant channels after filtering at 300-625Hz (Watson *et al.*, 2016). Each second was initially automatically categorized into WAKE, nonREM, REM and Microarousal (MA) states using a classifier based on a combination of spectrographic power and LFP-derived EMG tone. These sleep states were manually checked and corrected after initial automated scoring. All data are publically available for download at CRCNS: http://crcns.org/data-sets/fcx/fcx-1.

All analyses were carried out using a combination of custom code in MATLAB (Natick MA), the TStoolbox (https://github.com/PeyracheLab/TStoolbox), the FMA Toolbox (http://fmatoolbox.sourceforge.net/), and a Buzsaki laboratory code library: “buzcode” (https://github.com/buzsakilab/buzcode).

Power spectral analysis was done via Gabor wavelet convolution using MATLAB (awt_freqlist.m in buzcode collection). Wavelet bands were 50 logarithmically spaced bands between 1Hz and 625Hz (the Nyquist frequency for our signal based on our sampling rate of 1250Hz), but were rounded to unique integer values to prevent overlap in heuristically nearby frequencies. Wavelet data was sampled at 1250Hz, ie a 1:1 ratio with our LFP file, despite the fact that bins tended to be much larger than that interval.

Binwise analyses were done via creation of specific bins using TStoolbox that were then analyzed for either spike rate or spectral components or both. Phase-modulation analysis was carried out using circular statistics tools inside buzcode. For shuffled controls, spike times within a 3 cycle timespan (for whatever frequency is in question) were randomly shuffled with a flat probability density function. Correlation analyses were all carried out using Pearson correlations. For exploration of log value correlations for the final analyses presented, data were normalized to have a maximum of 1 and minimum of the minimum specifiable data above 0, in order to prevent taking log of negative values.

## RESULTS

### Total population firing activity correlates with gamma LFP power

We recorded a combination of LFP and action potential activity from 11 Long Evans rats over 27 sessions as they freely behaved in their home environments for at least 2 hours per recording. After spike detection and sorting using klustakwik, we separated units into putative excitatory (pE) or putative inhibitory (pI) types based on their waveforms (Stark *et al.*, 2013) and we classified each second of the recordings into WAKE, nonREM, REM or intermediate states (Watson *et al.*, 2016).

Our initial analysis was aimed at correlating combined population neuronal activity with broadband gamma power as has been seen in human data (Mukamel, 2005; Nir *et al.*, 2007, 2008; Ray *et al.*, 2008; Manning *et al.*, 2009; Miller *et al.*, 2014). The wideband (1 Hz - 20 kHz) data was low-pass filtered (< 1250Hz) and we performed wavelet decomposition at approximately logarithmically-spaced frequency intervals between 1 Hz and 625 Hz (see Methods). Based on previous work (Mukamel, 2005; Nir *et al.*, 2007, 2008; Ray *et al.*, 2008; Manning *et al.*, 2009; Miller *et al.*, 2014) as well as our own findings to be presented below, we defined a band of “broadband gamma” between 50- 180 Hz and correlated that against firing rates. Figure 1A shows z-scored firing rates of all pE units (n = 24) in 1-second bins from a single recording (green), all pI units (n = 5) from that same recording (red), integrated 50-180 Hz power in the same bins (blue) and EMG tone derived from the high-pass filtered LFP (black) in the same recording session during waking (WAKE), nonREM and REM sleep. Visual inspection of these curves suggests that firing rates correlate with gamma band LFP power. Both firing rates of individual neurons and broadband gamma (50-180 Hz) LFP power in one-second epochs varied extensively and showed an approximately Gaussian distribution on a logarithmic axis. We found positive correlations between summated population firing rates and broadband gamma power during Wake, nREM and REM for pE units and pI units; Figure 1C; Supplementary Figure 1).

**Figure 1.**
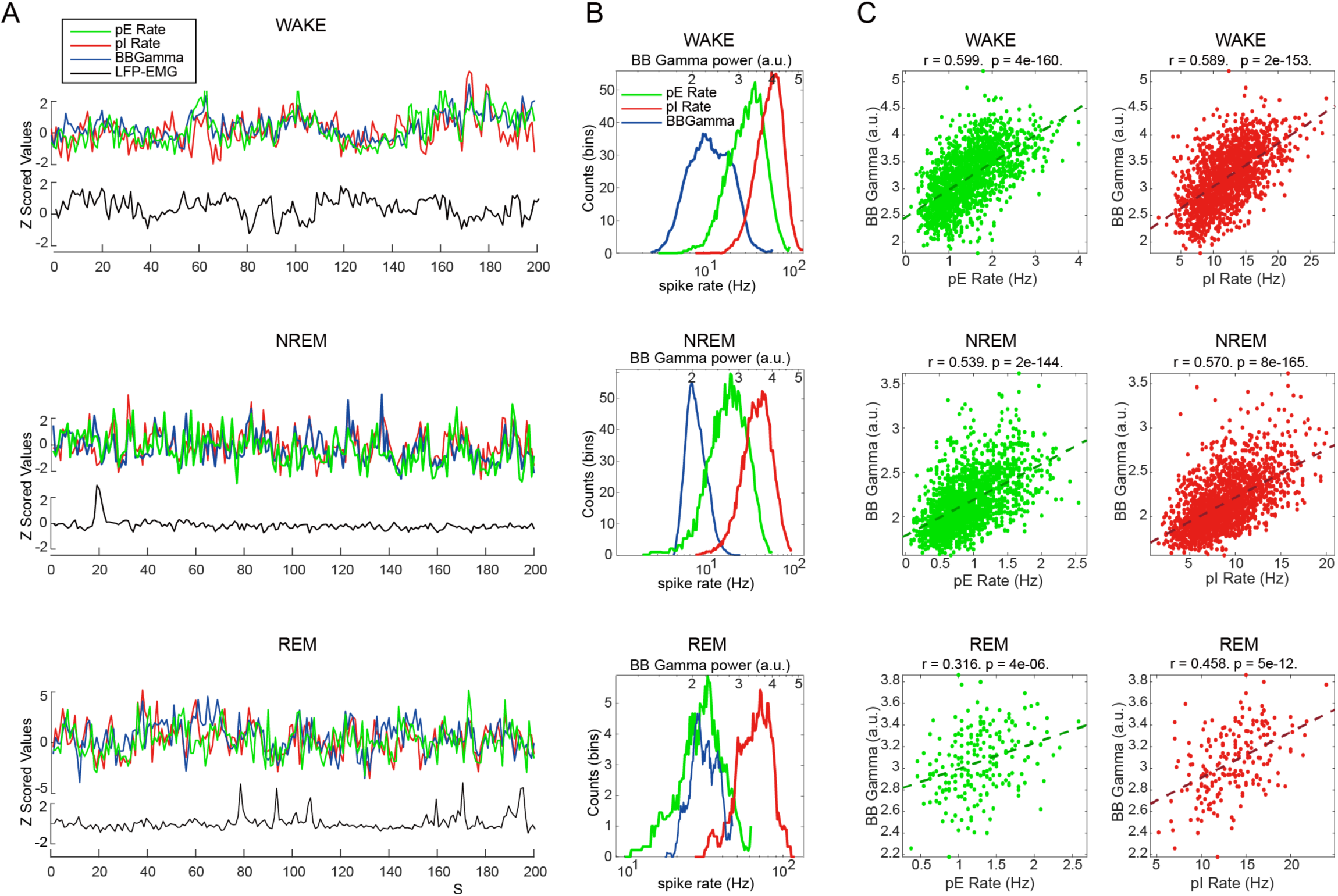
Correlation of population spike rate with broadband high gamma power. A. Broadband high gamma power (blue, 50-180Hz integrated power), summated spike rates of putative excitatory (pE) units (green), and summated spike rates of putative inhibitory (pI) populations (red) and LFP-derived electromyogram (EMG) (black) (see methods for extraction) plotted over time in various brain states and z-scored for comparison. Plots are all from one single recording binned at 1 second intervals for both LFP band power and spike rate and are separated by sleep/wake state: WAKE, NREM sleep and REM sleep. Same-state periods within a given recording were concatenated and here and two hundred seconds from each state are shown. B. Log-scale histograms of number of 1 second bins with each amount of Fourier power in the high gamma broad band range (blue), pE spike rate (green) and pI spike rate (red). Note that all of these statistics occupy their values in roughly log-normal distributions over time. Segregated by state. C. Bin-wise (1 second bins) scatter plots of correlations between broadband high gamma power (ordinate) and population spike rates (abscissa) in pE and pI cell populations across various states. Linear fits shown as dashed lines. Note Pearson’s r value of approximately 0.5 for population firing rate versus the broadband gamma across cell types and states (WAKE, NREM, REM).

### Single unit firing rate correlations with single frequency band powers in various brain states

Next, we quantified the Pearson correlation coefficients between the firing rate of each individual neuron and the powers of every frequency band analyzed in the local LFP at 1-second bin size in order to examine the full family of correlations between neurons and frequency bands in the cortex (Figure 2). We found that in the WAKE state both pE and pI neurons tended, on average, to be positively correlated with theta (4-8Hz) and gamma power (30-180Hz), but they tended to fire less when delta (1- 4Hz) and spindle-beta band (10-30Hz) power were increased (Figure 2A). A similar profile was seen in REM sleep. On the other hand, in nonREM sleep all bands above 5Hz were positively correlated and bands below 5 Hz were negatively correlated. In all states, the highest frequency bands (above 300 Hz) showed decreased ability to predict spike rates relative to the 50-180Hz band. We compared these power-spike rate curves, or “correlation spectra” with the LFP power spectrum in each state (Figure 2B) and observed that LFP spectral power qualitatively differed in profile from correlation spectra (LFP power presented corrected for 1/frequency decrement of power by multiplication of each value by the frequency of the band – for easier visual comparison against Figure 2A). This observation indicates that the overall curve of correlation spectra are not simple reflections of LFP power, indicating that they represent some specific process separate from LFP power alone. To test for significance, we shuffled the spike rate bin values versus the frequency band powers for each cell 200 times. The 95% confidence intervals are indicated as partially transparent shaded regions in figure 2A, we find that nearly all spike-power correlations are greater than expected by chance.

**Figure 2.**
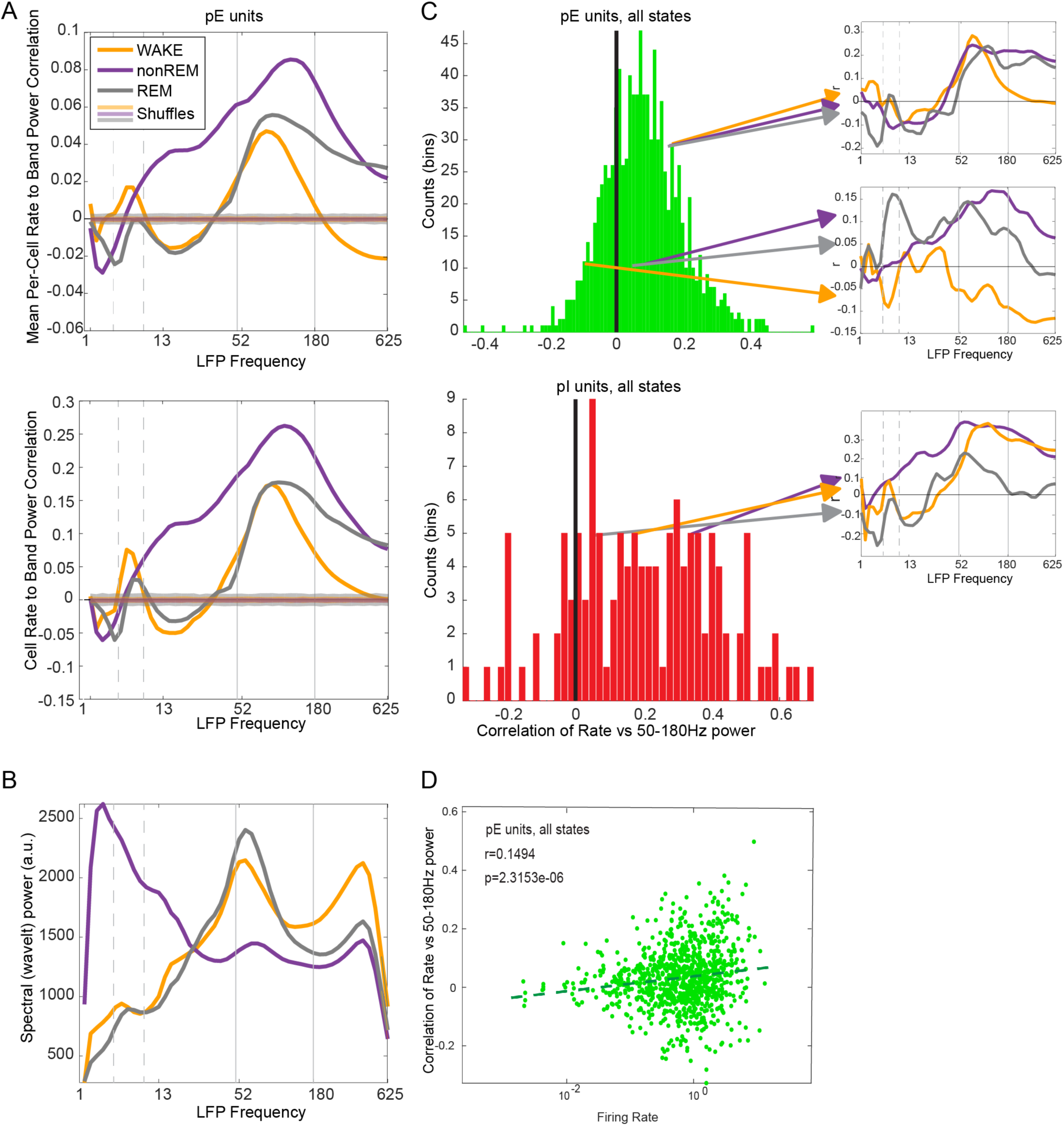
Variance in band power – spike rate correlation of two neuronal types with varying frequency bands. A. Means of Pearson correlation (r) values for individually-analyzed pE and pI unit firing rates at various LFP frequency bands. Units were analyzed and correlated individually and then averaged. Data was wavelet decomposed and both wavelet power within bands and spike rates were binned at 1 second intervals for comparison. WAKE, NREM and REM are shown in different colors on each plot and pE units are in the top plot, pI units in the bottom plot. Partially transparent lines with widths spanning the zero correlation line indicate the 95% confidence interval of 200 shuffles for each cell class and state. On average neurons show positive firing rate correlations with 50-180Hz oscillations across states, but NREM shows a wider band correlation spanning all frequencies except delta band (1-4Hz). Theta band power also tends to be positively correlated with spike rates in WAKE and REM. Note while in REM the highest frequency bands do not have positive correlations with spike rates, they do in WAKE. B. Spectral power in WAKE, NREM and REM states, corrected for 1/f trend by multiplication by f, to facilitate comparison against panel A. Band power-spike rate correlation values in each frequency band are not directly proportional to power in those frequency bands. C. Histogram of single-unit correlations with the 50- 180Hz band with 1 second bins. pE units, top, show a mean positive correlation but with a greater proportion showing negative correlations than pI units (bottom). Band-correlation plots similar to panel A for example single units are shown at right, top two are pE units and bottom is a pI unit. The topmost shows positive correlations with 50-180Hz band in all states (WAKE, NREM, REM) as shown by lines with same color code as in panel A, while the second unit from the top shows negative correlation with 50-180Hz power in WAKE but not in NREM or REM. Bottom plot is from a pI unit with positive correlations of spike rate vs 50-180Hz power in all states. D. Higher firing rate pE units show greater correlation with 50- 180Hz power.

In examining the distribution rather than the mean of individual neuronal correlations with the 50- 180 Hz broadband gamma (Figure 2C) we found that that while the spike rates of many pE neurons are positively correlated with power in this band, many also exhibit prominent negative correlations. This variance between neurons seems to account in part for the lower correlation values seen here compared to population-wise analyses (Mukamel et al, 2005 and Supplemental Figure 1). An additional component of the somewhat lower correlation values seen in many of these analyses may result from the particularly high correlations seen during delta-rich states, especially at time bins shorter than delta frequency (Supplemental Figure 1 and (Nir *et al.*, 2007; Mochol *et al.*, 2015).

Additionally, many neurons flipped the direction of their modulation by 50-180 Hz band power across brain states as can be seen in the examples highlighted at right in Figure 2C. Supplementary Figure 2 shows an analysis of consistency across states for this broadband gamma-spike rate correlation for each cell in the dataset. We found that switching of firing rate correlation with broadband gamma as a function of brain state is common across many pE and pI units.

### Contribution of spike waveforms and EMG to high frequency LFP power

While wavelet power spectra during WAKE in many sessions showed a recognizable peak at approximately 400 Hz (Figure 2B), no such peaks were observed in the spike–LFP power correlograms. We assumed that the highest frequency band is particularly impacted by contamination by both muscular activity and spiking events themselves. EMG was measured using the Pearson correlation across physically distant brain electrodes in the 300-650 Hz range, as this has been found to correlate well with muscularly-measured EMG activity (Schomburg *et al.*, 2014). We examined the impact of EMG activity on the spectrum, which can be seen even by eye on the spectrogram (Figure 3A) by subdividing our WAKE episodes into 1-second-long bins and ranked those into groups depending on the relative amount of EMG activity. The periods of increased EMG power correlated with increased power above bands from approximately 50 Hz up to 650 Hz (Figure 3B). Strikingly, REM sleep epochs showed the lowest power in these high frequency bands. This seems to relate with the known drop in muscular tone during REM and adds further support to the contribution of EMG activity to spectral LFP power during WAKE.

**Figure 3.**
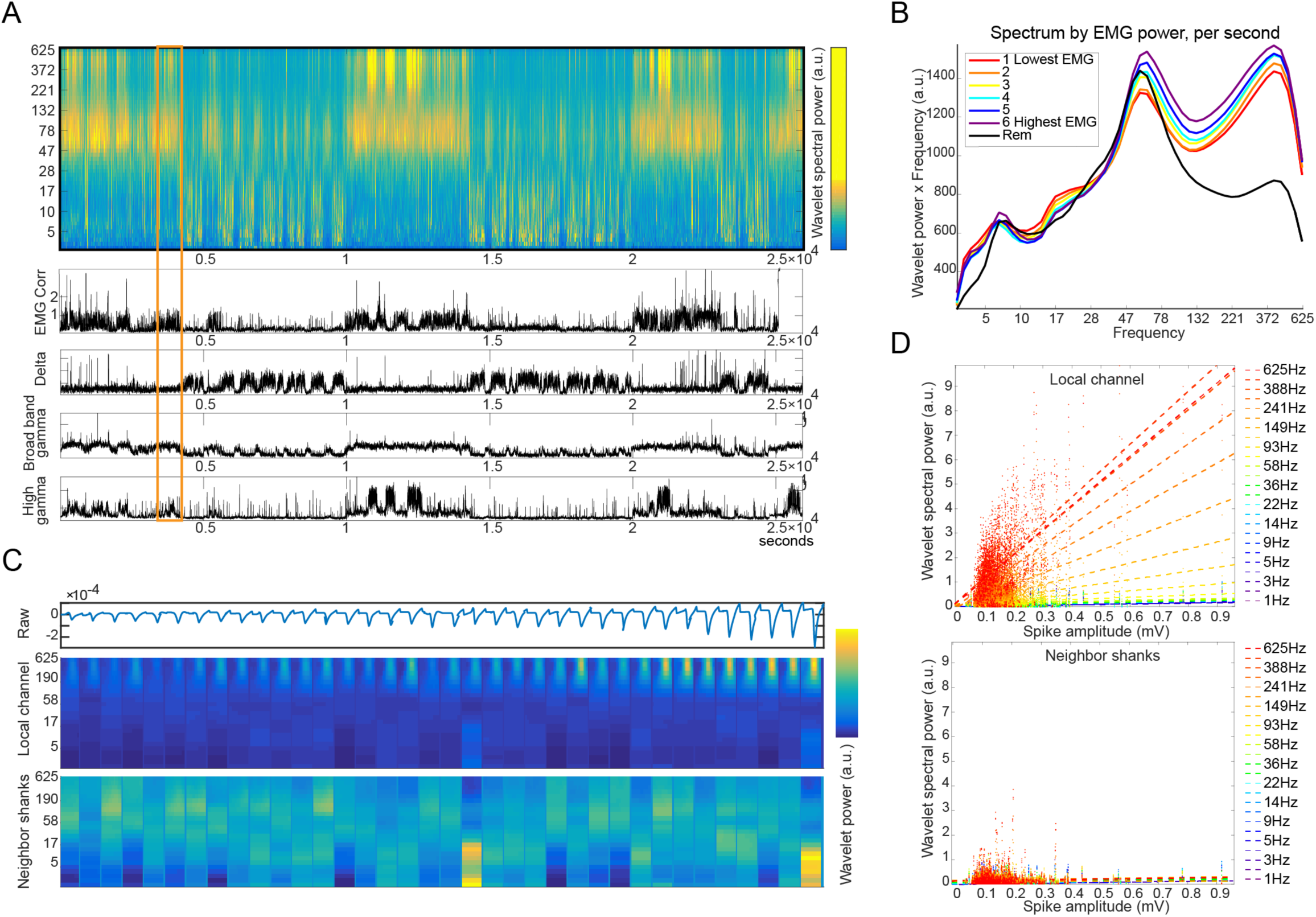
Electromyographic and spiking contributions to LFP signal. A. Example wavelet spectrogram and EMG recording from a single session. Power in various (but not all) frequency bands are shown in line plots below spectrogram. Note general tendency for spectral power in the 300-600Hz range to increase during period of high EMG tone. B. EMG state impacts LFP: WAKE epochs (rainbow color lines) with higher EMG tone (purple) have higher power in the high frequency range than low-EMG seconds (red). REM is shown in gray and shows relatively decreased power in the highest frequency bands. C. Spikes correlate with instantaneous power in the highest frequency band and larger spikes have more such impact. Panel shows data from a single example session wherein spike waveforms are presented sorted from smallest to largest at top and below is shown the impact of each spike above on the power spectrum of the local channel. More distant channels (bottom most) residing at 200um away from the channel with the spike show diminished high frequency power during spiking. D. Dataset-wide quantification of the effects of spiking on temporally locked spectral power. Top panel shows the correlation of spike size to the spectral impact over various frequency bands examined on the channel with the spike itself. The red dots and fit line for those dots relate to the highest frequencies and bluer tones relate to lower frequencies. Note that power in higher frequencies is strongly dependent on spike size, while this effect is diminished in lower frequencies. Bottom panel shows the same analysis but in a case wherein the LFP originated from neighbor shanks 200um away from the unit spike. Note the reduced impact on this LFP by distant spikes.

Spike waveforms have also been shown to impact LFP in high frequency bands in the hippocampus, especially during strongly synchronous neuronal firing events (Ray & Maunsell, 2011; Belluscio *et al.*, 2012; Schomburg *et al.*, 2012). We quantified this effect on the cortical LFP using spike-sampled averages and we then dissected that analysis based on spike amplitude and distance from electrode (using the known geometry of the implanted probes). For every recording session, we calculated the average raw spike waveform for each identified unit and then ranked those spikes from lowest to highest amplitude (Figure 3C). We then calculated the LFP impact of these raw waveforms and sorted them by the size of the initial raw waveform. Based on the increased intensity (yellow) at the highest frequencies of the largest spikes in this plot, it is visually evident that the highest frequency bands are increasingly impacted by the largest amplitude spikes. We quantified this effect by demonstrating increasing slopes and correlation coefficients of spike size-spectral power with increasing spectral bands (Figure 3D) and found statistically significant Pearson correlations between raw waveform size and LFP wavelet spectral power for all frequency bands increasing from from p = 10^−6^ for the 1Hz band to p = 10^−92^ for 625Hz band. The significant correlations in the high frequency bands disappeared when we compared the impact of a spike on the LFP of a channel 200µm away (since the shanks of our silicon probes are spaced 200µm apart), although positive correlations remained in the range of p = 10^−10^ for bands between 1 and 36Hz. Previous research has shown that spike amplitude decreases rapidly with distance and at lateral distances > 150 µm, the amplitude is near the biological noise (Henze *et al.*, 2000). Yet, since many neurons are included in this annulus, even their very small amplitude contributions can add significantly to LFP power (Schomburg *et al.*, 2012), we also compared LFP power > 200 µm (two or more shanks away) from the recorded spike and found even less contribution at those distances (Supplementary Figure 3).

### Timescale of correlation of spike rate to LFP power

The above analysis used 1 second bins and was not able to determine the time precision of the correlation between LFP power and spike rate. To address this issue, we varied bin sizes in our LFP band-spike rate correlations. Figure 4 shows averages of correlation values from all pE and pI neurons in the dataset across all frequency bands at multiple time bin sizes (Figure 4A). In general, we found that across most bin sizes and sleep/WAKE states, larger bin sizes lead to greater correlations between spike rate and LFP power - up to a peak of around 100 second bins. Bin sizes greater than 200 seconds could not be consistently assessed given brain state switches, especially for REM. The 50-180 Hz band continued to stand out with the highest positive correlation to spike rate, followed by the theta band during WAKE.

**Figure 4.**
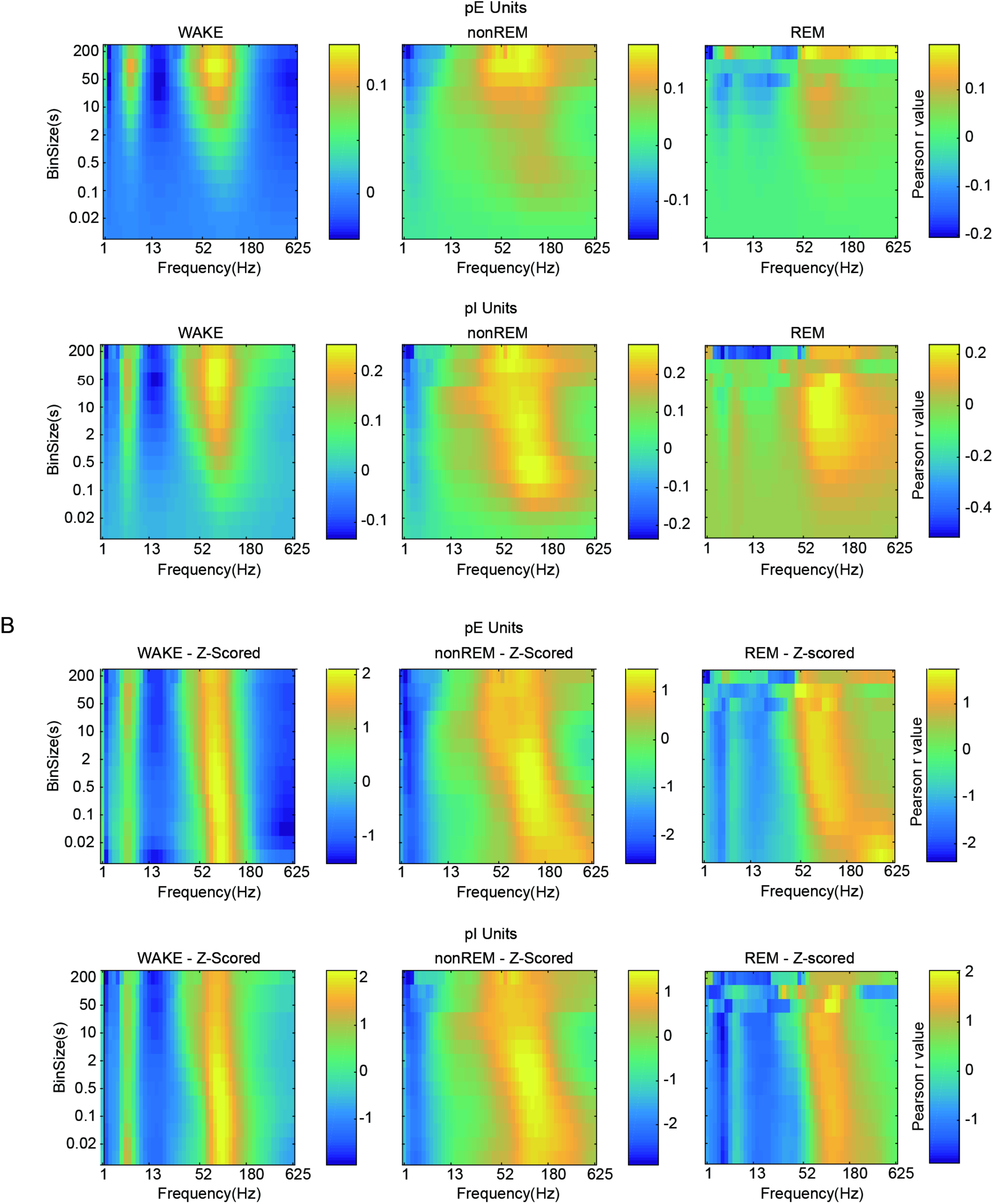
Time bin size versus frequency band dependence of spike-rate to band-power correlation. A. Displays of Pearson correlation value in color (see color bars) averaged over all cells in the dataset for each frequency band (ordinate) for various bin sizes (abscissa, identical bins used for both spike rates and frequency band power quantifications) for pE and pI units. Units were analyzed and correlated individually and then averaged. Pearson r values across WAKE, NREM and REM states are shown. Across states, maximum correlation averages are seen around 50-180Hz in bins of 50-200 second sizes. B. Pearson r values z-scored within each bin size in order to show relative preference within each bin size for the various frequency bands. Maximal population-wide average is now seen around the 50-180Hz frequency band at around 0.1-2 second bin size. Smaller bin sizes correlate best with slightly up-shifted broadband gamma frequencies, though not in proportion to the reciprocal of the oscillation frequency and not to the point where the highest frequency band shows peak power.

It is possible that this increase in correlation values with increasing bin durations could relate to improved sampling of a noisy process by longer bins. Under this assumption and in order to extract frequency band preference within each bin width, we z-scored the values across frequency bands within each bin size (Figure 4B). Across all bin sizes, the 50-180 Hz band continued to be most positively correlated with firing rates. Interestingly, smaller bin sizes correlate with a slight increase in the preferred frequency band for spiking rates, though these increases were not proportional to the frequency associated with the reciprocal bin size itself. Within this z-scored series of plots, is a peak in the range of 0.1 to 2 second wide bins for the 50-180Hz band, meaning that these may all be reasonably equivalent bin sizes for experimenters to choose as surrogates for spike rates.

We wondered what single combinations of bins and bands individual neurons most showed preference for. To do this, we then used this z-scored data to search for peaks for each neuron across bin sizes and frequency bands (Supplementary Figure 4). These two dimensional histograms indicate that individual unit spike rates showed most preferred bands and bin sizes with distributions similar to the amplitude of the mean r value plots shown in Figure 4B – with peaks at approximately 90 Hz LFP and 1-second bins with small variance across brain states. Despite this relationship, overall preferences for gamma band power as a predictor of spike rates, considerable diversity exists with many units firing rates being best predicted by low frequency bands.

### LFP phase modulation of spikes

While spike rate correlations to LFP power represents one relationship between spikes and LFP, we wondered if the very different metric of spike timing locked to certain phases of an oscillation might show us similar spectral profiles to our rate-power metric. To quantify the spike-LFP phase relationship in depth, we measured phase-modulation of spike timing to specific phases of the LFP at all frequency bands across all units using the mean resultant length (MRL) of the vector to measure the phase of firing of spikes for each unit. We found that both cortical pE units and pI showed phase modulation at many frequency bands (Figure 5A-B), and that this modulation was greater than expected by chance, as compared to spike timing-shuffled LFP. Excitatory neurons lead the inhibitory neurons by approximately 25 degrees in phase space (Supplementary Figure 5D,E), which corresponds to a 0.4-1.5 ms delay depending on the oscillation frequency, and this delay is consistent with models that gamma oscillations are mainly brought about by a E-I recurrent mechanisms (Csicsvari *et al.*, 2003; Moca *et al.*, 2011; Buzsáki & Wang, 2012).

**Figure 5.**
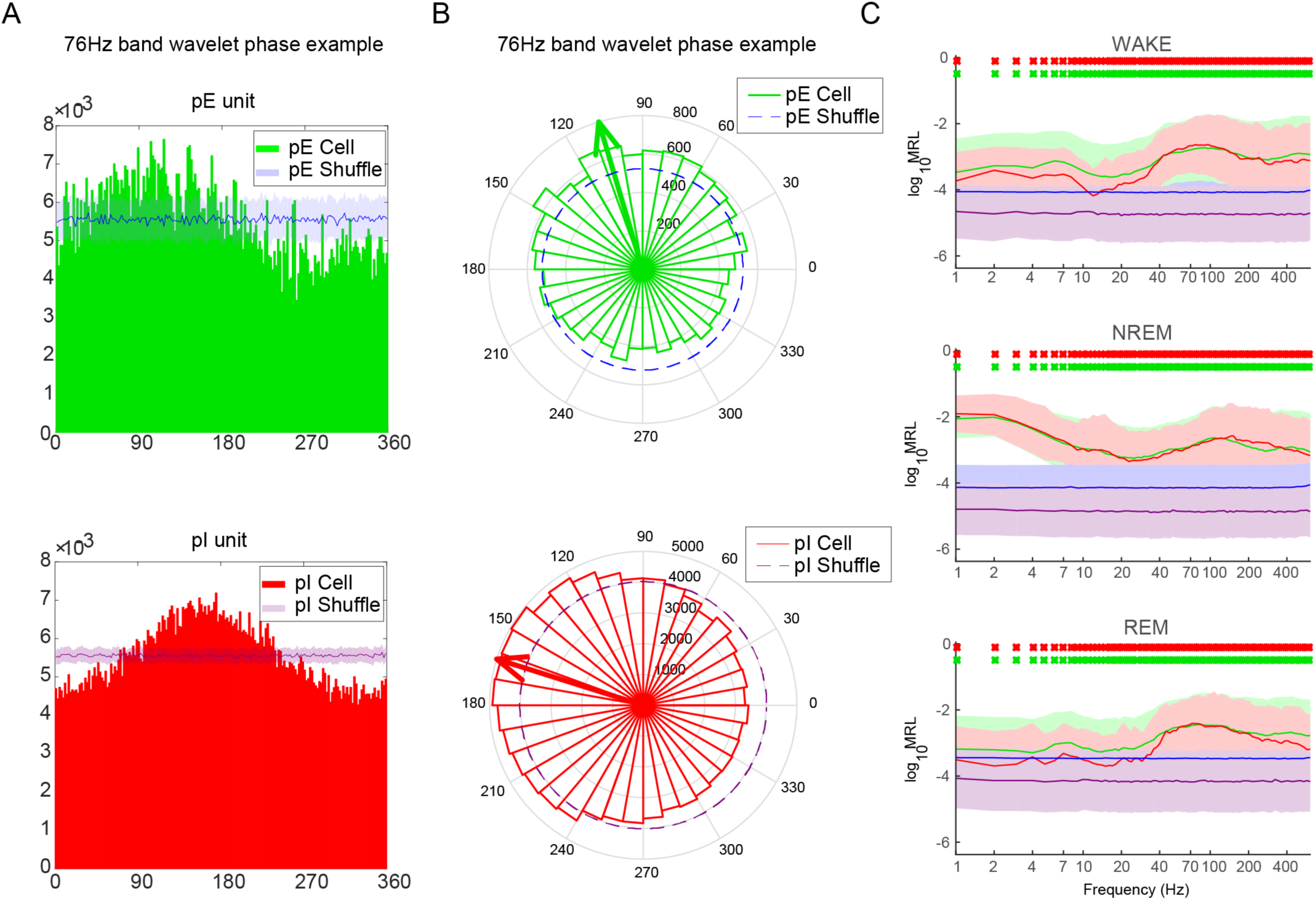
Coupling of spikes to LFP phase - by frequency band and by state. A. Phase modulation of an example pE unit (top panel) and a pI unit (bottom panel) by 76Hz frequency phase during high-power periods. Unit spikes are shown in green (pE) and red (pI), and modulation values found in 200 shuffled datasets shown in blue (top) and purple (bottom). Shaded regions surrounding central tendency lines indicate the standard deviation over neurons for both real and shuffled data. Shuffling was done by randomizing the timing of each spike over a local interval representing three times the cycle duration for the frequency being analyzed. B. Rose plot and mean resultant vector for same examples used in panel A. Mean resultant vector shown as vector arrow and mean resultant length (MRL) of this vector will be used as a generalized quantification of modulation of spike generation by LFP phase. C. Log of mean resultant vector lengths averaged across all pE units and pI units in the dataset, across all frequencies, stratified by state. Significant increases in MRL relative to shuffled datasets are shown in red and green x’s above each bin size, as assessed by a 1-sided t-test with Bonferroni correction and alpha set to p = 0.01. All frequencies showed modulation above chance for their respective cell type.

We then used this MRL metric (‘preferred phase’; (Csicsvari *et al.*, 1999)), averaged over all pE neurons and pI units across and all frequency bands in different brain states, and we again found spikes were best modulated by the 50-180 Hz broadband gamma range during WAKE and REM, while during nonREM lower frequency bands also showed more positive phase modulation than 200 shuffled datasets (Figure 5C). These phase modulation depths profiles were similar to the rate-power profiles shown in earlier figures. Significant increases in MRL relative to shuffled datasets were assessed by a right-tailed Wilcoxon rank sum test with alpha set at p = 0.01 after Bonferroni correction. All frequencies showed modulation above chance for their respective cell type.

To further assess the relationship between spiking and LFP, we compared the relative influence of high LFP power versus low LFP power epochs of each oscillation frequency (Supplementary Figure 5). With this analysis we found that high-power epochs of high-gamma band again tended to show a preferential prediction of neuronal spike phase, especially during REM sleep and WAKE, although other frequency bands also showed a power-dependent phase modulation of spiking. During nonREM sleep, delta-band oscillations in the 1-5 Hz range exerted a particularly power-dependent correlation with spike phase. In the low LFP power group, significant phase-modulation of spikes occurred only in the >100 Hz range, suggesting that in the very high frequency range, the spike-LFP correlations may reflect spike coupling to spike afterpotentials themselves (Penttonen *et al.*, 1998; Belluscio *et al.*, 2012; Buzsáki *et al.*, 2012; Zanos *et al.*, 2012) rather than to genuine network oscillations.

### Peri-spiking field potentials and population spiking

Putative excitatory and inhibitory units were similar in most respects as observed in our analyses thus far using LFP as the independent variable. In the following analyses, we regarded spikes as the independent variable and constructed spike triggered averages of wavelet power (Figure 6). Wavelet spectra were calculated from the LFP recording from the neighboring shank (200 µm away) from the recorded spike to minimize spike contamination (Figure 3). Wavelet spectra triggered by pE spikes showed an increased gain of 70-150Hz band power after the spike, with the low frequencies lasting up to 200 ms after the pE spike. In contrast, after pI spikes the gamma activity increase was more transient (< 100 ms) and no increase of > 200 Hz power was present around the pI spike (Figure 6; Supplementary Figure 3). Theta band power was typically increased prior to both pE and pI spikes.

**Figure 6.**
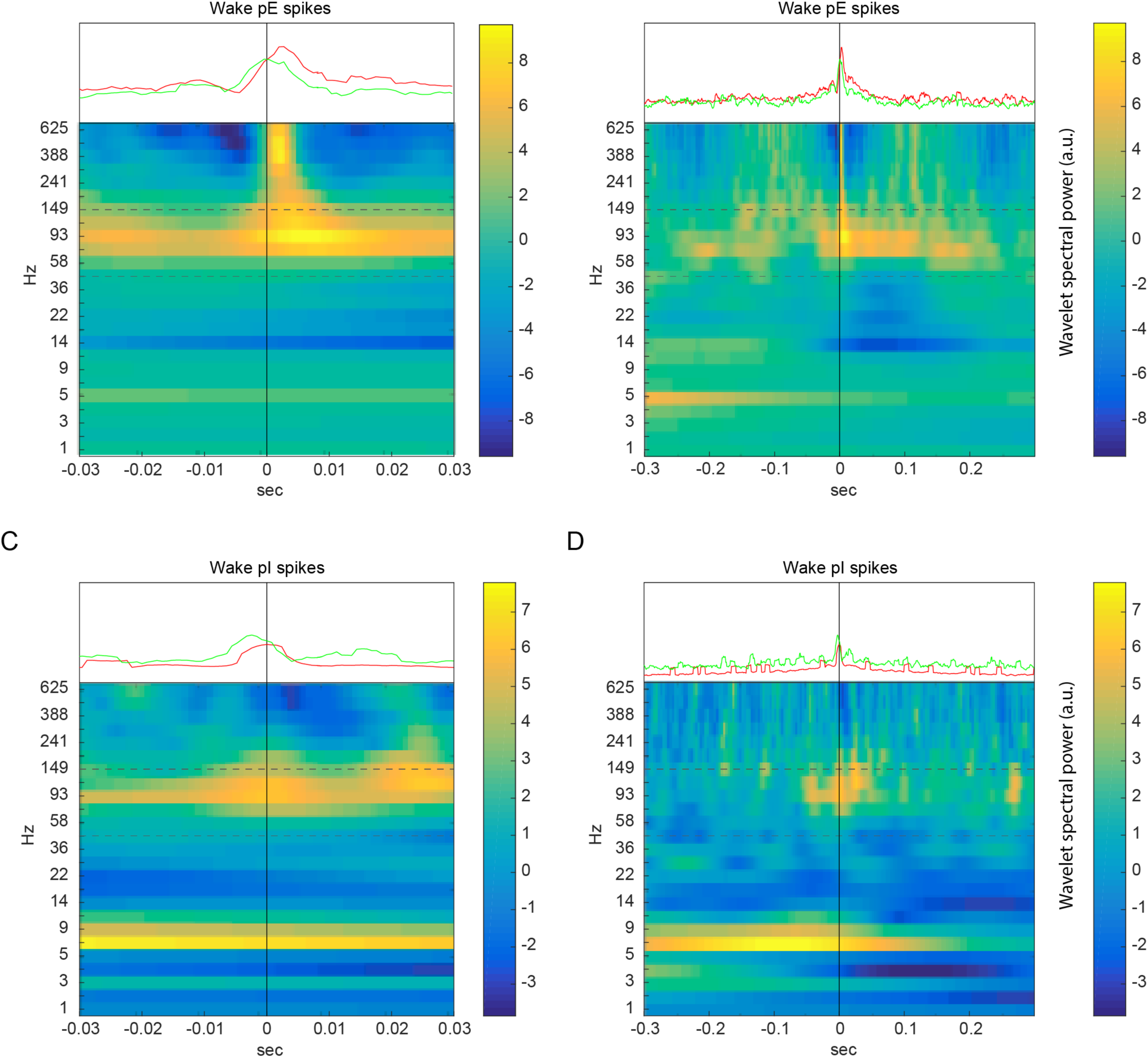
Spike triggered spectrograms for pE and pI units. A and B show spike triggered spectrograms for pE units at ±30ms (A) and ±300ms (B). LFP channel for wavelet spectra is from shanks neighboring the unit shank (200um away). C and D show spike triggered spectrograms for pI units at ±30ms (C) and ±300ms (D), from neighboring shanks. 50-180Hz power is increased around the time of spiking of both pE and pI units, with the up-modulation of power somewhat shorter around pI spikes. After pE unit spiking, 12-45Hz power is decreased and 300-625Hz power is rhythmically modulated at approximately 4Hz. Green and red lines above each plot show spike-triggered firing of the surrounding pE and pI populations recorded simultaneously with (but not on the same shank as) the trigger unit. Note pE units on average fire 3ms before pI units.

In addition to spike triggered wavelets we also gathered spike-triggered firing rates of concomitantly recorded pE and pI units (excluding from the same shank as the trigger unit due to difficulties detecting spikes within short latencies on the same shank). We found that pE units were roughly synchronous with each other with a cross-correlation width of around 10ms, whereas pI units tended to follow pE units by approximately 3ms on average (Figure 6A and 6C), complementing the pE-before-pI findings of the preferred phase of spikes to the gamma cycle (see Supplementary Fig. 6 for similar analysis in nonREM and REM states).

### Excitatory-inhibitory network balance and LFP

To determine how excitatory-inhibitory balance may relate to LFP oscillations, we calculated the pE unit to pI unit population activity ratio for each second (E/I ratio), and generated LFP spectra for each second based which of ten E/I ratio bins it fell into. To calculate E/I ratio we counted the total pE unit spikes (across all units in a given session) in a given time bin as a proportion of all pE+pI spikes and then z-scored this value for each recording in order to compare across recordings. We categorized each second of each recording into one of ten E/I ratio bins and accumulated average LFP wavelet power spectra from the seconds that match each of those bins (Figure 7). We found that the magnitude of E/I ratio correlated with LFP band power, although in a complex manner. Similar to the firing rate versus 50-180Hz band correlations (Figure 2), we found that the E/I ratio also correlated with the power in the 50-180Hz band with lower E/I ratios linked to higher LFP power in the broadband gamma band. This was the case not only in the z-scored data (Figure 7, dotted bands) but also in the raw spectra (Supplementary Figure 7). In addition, we broke the WAKE state down into epochs of high movement for at least 5 seconds (WAKE-Move5s) or low mobility for at least 5 seconds (WAKE- Nonmove5s). The two wake-related states without movement: REM and WAKE-Nonmove5s showed similar patterns of modulation of the power spectrum by E/I ratio wherein high frequency power decreased as E/I ratio increased and low frequency (5-30Hz) power was oppositely modulated co-modulated with high frequency bands in REM. This may relate to the tendency for synchronous delta-power dependent states to occur during quiet wakefulness – wherein nonREM-like comodulation of 1-30Hz power is frequently seen (Watson *et al.*, 2016). During nonREM, power in all frequencies was negatively modulated by E/I ratio. Many of these variances in frequency band modulation can be understood from large differences in low frequency power in the nonREM and REM states in the raw spectra (Supplementary Figure 7). Regardless, the modulation of the 50-180Hz band downward with increasing E/I ratio is common across all states and all analyses.

**Figure 7.**
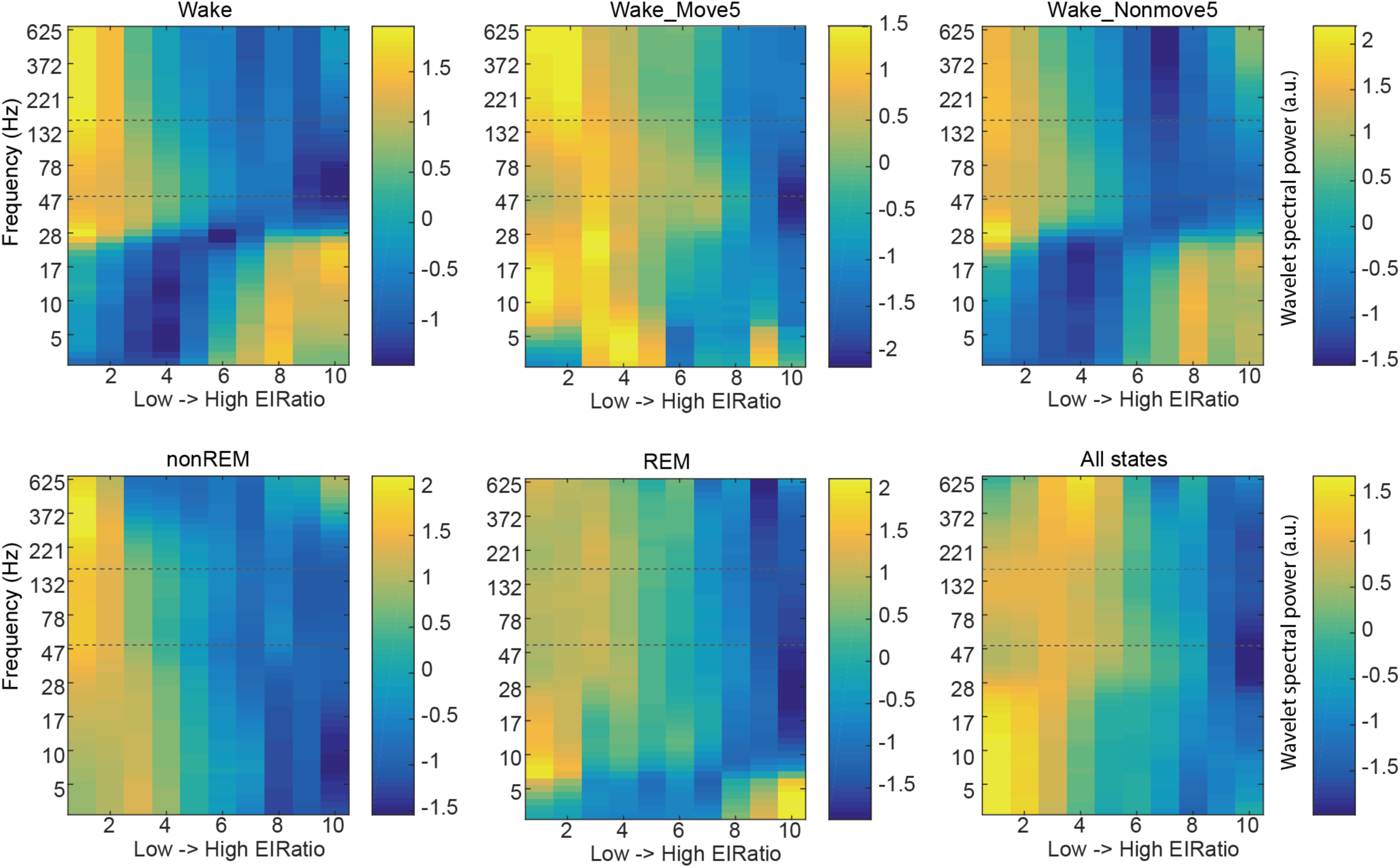
Modulation of LFP by second-wise excitatory-inhibitory state of the network. Recordings were binned at 1 second and in each bin the total E and I unit spikes were calculated and used to rank the bins by relative excitatory-inhibitory ratio (pE spikes / (pE spikes + pI spikes)). Bins were ranked into 10 groups and average wavelet spectra were calculated for each group and averaged across all recordings in the dataset. Shown are averaged binned spectra as vertical stripes, data was smoothed between bands over a 3 band width and then z-scored within each frequency band in each brain state. The 50-180Hz band is highlighted in each plot using dotted horizontal lines and is consistently down-modulated as EI ratio increases while lower frequencies show more varied modulation by EI ratio depending on the state.

### Wide band correlation-based prediction of spike rates

A final analysis was to find best LFP-based predictors of population firing. Based on our analyses, we saw that while the 50-180Hz band did indeed strongly correlate with population firing, we wondered if we could use information available from other frequencies to improve our LFP-spike rate predictability. In order to more broadly search for predictors of spiking, we correlated various LFP metrics against summated population spike rates in each bin, all within the WAKE state. Our goal was to explore the predictors beyond band-wise power as had been used in prior studies but, as a comparison we did perform band-wise powers with spikes in various bands, including delta, theta, sigma, beta, low gamma, broadband gamma, epsilon 200-600Hz power and 1-200Hz power.

Next, we used dot product-based projections of within-bin spectra against various frequency spectra against including WAKE, WAKE-Move5s, WAKE-Nonmove5s, nonREM, REM, microarousals, and WAKE minus nonREM. To avoid bias for low frequency predictors based on the known 1/frequency-based decrement of oscillatory power, both predictor and test spectra were whitened via multiplication of the power value by the frequency.

Finally we defined “correlation spectra” based on band-wise correlations with firing rates as shown in Figure 2A and we projected those against the spectra in each temporal bin. Separate correlation spectra were created for both pE and pI unit types from various brain states, again as shown in Figure 2A. Since within-recording correlation spectra would not be measurable in circumstances without unit isolation, as is the case for many experimenters, the correlation spectra used for our analyses were the dataset-wide mean correlation spectra with the idea that these can be used as predictors for any recording going into the future.

Figure 8 shows histograms of correlation r values per recording for a subset of the predictors we used (n = 27 recordings over 11 animals). The projection against the correlation spectrum had more consistently positive correlation against pE population spiking, with higher mean, median and minimum correlations, than other methods used here. The complete set of analyses with all predictors included are shown in Supplementary Figure 8, again with the projection against the WAKE-based correlation spectrum showing more reliably high correlation values than other predictors. The profile of predictor correlations for pI cells was generally similar to that of pE cells while log-transformed predictors tended to worsen correlation values and log-transformation of spike rate histogram values did not qualitatively change correlation values (not shown).

**Figure 8.**
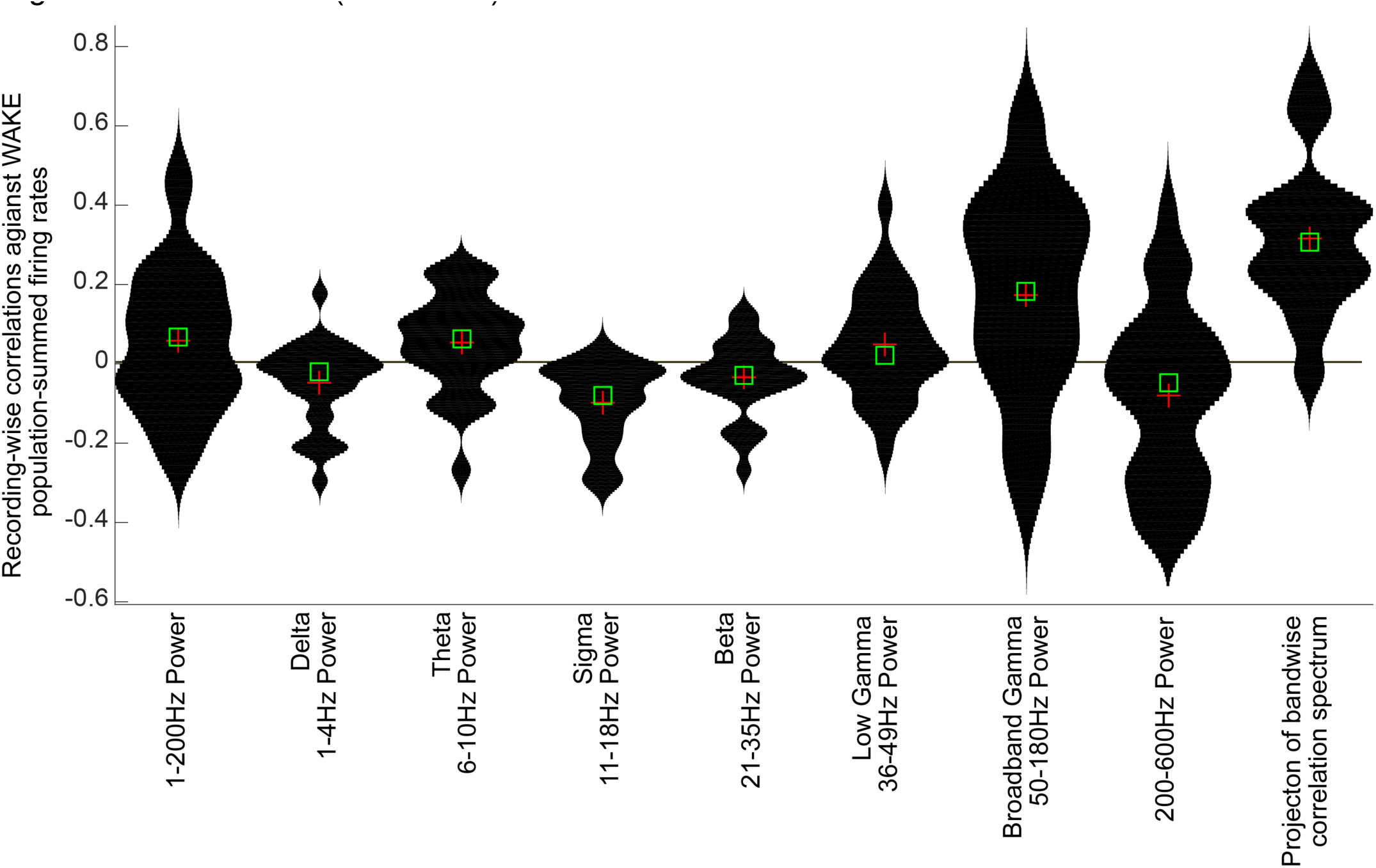
The correlation spectrum predicts overall spiking better than frequency band powers. Shown are recording-wise Pearson correlation values between summated population spike rates of pE units and with LFP predictors in the same bins. Violin plots represent distributions of correlation values over recordings for each metric. Mean values for each metric shown with red crosses, medians shown as green squares. The first 8 columns show spike-LFP correlates based on power in various power spectral bands as indicated, including the 50-180Hz broadband gamma power used elsewhere. All spectra are corrected for the general 1/frequency falloff in power via multiplication of power values by the frequency itself, in order to not bias for low frequency predictors. The last column is a dot-product projection of the bin-wise power spectrum against the correlation spectrum (orange curve shown in upper panel of Figure 2A). This yields correlations with higher mean and median powers and a higher minimum value than other methods examined.

## DISCUSSION

Using high-density recordings with high unit yields in rats, we have shown here that LFP power in the 50-180Hz band correlates positively with increased unit population firing rates, supporting previous observations (Mukamel, 2005; Nir *et al.*, 2007, 2008; Ray *et al.*, 2008; Manning *et al.*, 2009; Miller *et al.*, 2014). We report that other frequency bands also carry both positive and negative predictive information about firing rates (Kayser *et al.*, 2007). Additionally, we show that correlation is variable across sleep-wake states, with large variability across individual neurons and that the timescale of spike-LFP correlation is highest in the range of 100 – 2000 milliseconds. We validated these measurements using LFP phase-modulation of spiking of cortical neurons. We also show that power in the gamma to high-gamma range during waking states correlates with lower excitatory-inhibitory ratio. Finally, we provide a new metric, dot product projection of the correlation spectrum, for quantifying the relationship between spiking and LFP, and show that it provides stronger correlations values than previous methods.

### Spike-LFP coupling can arise from various mechanisms

Spikes can relate to LFP in multiple ways and a full understanding of spike-LFP coupling should take them all into consideration. First, in the simplest case, measured oscillations can result directly from transmission of upstream oscillations – either regular or irregular. More specifically, synchronous afferent drive can bring about synchronous depolarization of the target dendritic region, often coupled with feed-forward excitation of local inhibitory interneurons. Whether the target neurons are phase-locked to the projected afferent LFP pattern depends on multiple factors, most importantly on the frequency of the input. Frequencies up to approximately 80 Hz can be transferred, whereas above this frequency spike coupling with afferent drive is rare (Vaidya & Johnston, 2013; Buzsáki & Schomburg, 2015). Yet, spatially resolved gamma power in identified dendritic regions can provide important information about the magnitude of upstream drive (Colgin *et al.*, 2009; Lindén *et al.*, 2011; Berényi *et al.*, 2014; Schomburg *et al.*, 2014; Butler *et al.*, 2017; Fernandez-Ruiz *et al.*, 2017). A second mechanism for induction of oscillations is that irregular patterns can be induced by local circuit interactions. In this case, local neurons themselves bring about EPSPs and IPSPs in their local partners and, in turn induce transmembrane currents that can be recorded extracellularly in the form of LFP (Buzsáki *et al.*, 2012). Naturally, there is a correlation between spikes of the participating neurons and the PSPs they bring about. Thirdly, calcium spikes and other types of plateau potentials also produce transmembrane currents, thus contribute to LFP. Fourth, Na^+^ spikes of neurons working coherently or in an asynchronous regime have afterpotentials lasting 2-20 ms (Nadasdy *et al.*, 1998; Fernandez *et al.*, 2005), and such afterpotentials contribute power to the gamma frequency band. Finally, there are artefactual (ie not based on local transmembrane) sources of LFP, of which EMG contributes most power to the gamma band. Separation of these distinct mechanisms and quantifying their contribution to the compound LFP requires prior knowledge of anatomical connectivity combined with high spatial resolution methods and simultaneous recording of spikes from individual neurons (Buzsáki & Schomburg, 2015).

In addition to the multiple sources of LFP itself, the concept of spike-LFP coupling has at least two definitions. In the case of oscillatory LFP, a widely used measure is spike-phase coupling (‘spike-LFP phase’ coupling). Spike-phase coupling can be characterized by either establishing preferred phase (mean phase) of spikes to a particularly rhythm or the magnitude of phase-modulation (mean resultant length or MRL)(Csicsvari *et al.*, 1999). Another way of measuring spike-LFP coupling is to correlate spike density and LFP power in predetermined time windows (‘spike rate-LFP power’ coupling). These different measures can be affected by various factors, such as brain state, and may or may not correlate with each other. Here we used both metrics in search of convergent findings of co-modulation of spikes and LFP.

### LFP-spike coupling – comparison with previous works

Important differences in our experiments compared to those in humans should be noted and taken into account. Our data were obtained from the deep layers of the neocortex, whereas ECoG recordings largely reflect activity of transmembrane currents from the apical dendrites of layer II-V neurons. The rodent hippocampus generates large amplitude and highly regular theta oscillations (Vanderwolf, 1969), whereas theta activity in humans is more intermittent and of lower amplitude (Kahana *et al.*, 2001). While theta power in the LFP could reflect volume conducted fields, our finding of theta power-related unit firing in the neocortex and theta-phase modulation of neocortical units (Sirota *et al.*, 2008) indicates long-range communication between hippocampus and neocortex, which may be less expressed in primates. Despite differences in such details, we believe that our main findings in rodents generalize to the human brain as well.

### Spike contribution to the LFP

LFP data can serve as an extremely useful mesoscopic measurement of neuronal cooperation, with gamma band activity providing signatures of local processing within neural circuits. Human cortical surface recordings (electrocorticogram, ECoG), often performed in patients with epilepsy, do not give direct access to spiking information and so they rely upon LFP-related signals. Yet, it has been repeatedly suggested that higher frequency power largely reflects summed spikes of numerous neurons (Ray *et al.*, 2008; Schomburg *et al.*, 2012). In support of these suggestions, we found that LFP activity in the range 50-180 Hz was positively correlated with population firing rates and even firing rates of most individual neurons. This frequency range, showed the greatest power correlation (positive or negative) with spiking of any band. Furthermore, we also show that activity in higher frequency bands can often carry contamination from the spikes themselves (Figure 3). Spikes and their afterpotentials contributed to power in bands as low as 50 Hz (see also in (Belluscio *et al.*, 2012)), down to the band where true relatively narrowband network oscillations dominate (McAfee *et al.*, 2017; Saleem *et al.*, 2017).

It remains unclear why or how cortical neurons seem to fire more coincidently with increased power in the high gamma range. Does high gamma reflect a *bona fide* oscillation or simply spikes and spike afterpotentials (Belluscio *et al.*, 2012; Schomburg *et al.*, 2012)? Additional methods, such as simultaneous recording from several input structures, high-density recordings across multiple layers, current source density techniques, independent component analysis, de-spiking, cross-frequency coupling analyses and spike-LFP phase-locking, are needed to effectively separate LFP components that arise from synaptic currents and spike afterpotentials, respectively (Csicsvari *et al.*, 2003; Canolty *et al.*, 2006; Sirota *et al.*, 2008; Zanos *et al.*, 2011; Belluscio *et al.*, 2012; Fernandez-Ruiz *et al.*, 2012; Schomburg *et al.*, 2014; Buzsáki & Schomburg, 2015). Furthermore, appropriate methods are needed to determine whether gamma power reflects an oscillation or results from summation of non-rhythmic events with time constants similar to the gamma waves (Mureşan *et al.*, 2008; Buzsáki & Schomburg, 2015).

### Variable LFP-spike coupling across different bands, states and cells

To establish a disciplined way of quantifying spike-LFP power coupling, we performed band-by-band correlational analysis at 50 log-spaced intervals between 1 Hz and 625 Hz and used the r values of the correlations of each of these bands (Figure 2A) to create a prediction vector for the optimal LFP-spiking correlation. By projecting the LFP from each of the time bins of interest against this vector, we were able to predict spike rates more reliably than with broadband high-gamma alone (Nir *et al.*, 2007), broadband LFP (1-200Hz, (Miller *et al.*, 2014)) or other specified frequency bands (Figure 8). Our unbiased method could find the most reliable preferred LFP band for the spikes of individual neurons. These findings revealed that while the majority of pyramidal cells had a positive correlation with gamma band power, a significant minority of neurons had negative correlations with gamma power.

Putative excitatory and inhibitory units had largely similar -spike rate correspondences although pE units showed more variance. This increased variance may largely relate to the slow oscillation of nonREM sleep because the difference between pE and pI neurons disappeared with bin sizes larger than 1s (corresponding to slow oscillations).

Within the pE units, the higher firing rate units appeared to have stronger correlation with broadband gamma power. It is not clear whether this observation is due simply to spike-LFP contamination, statistical features or whether it is a true physiologic difference between cell types, although it has been observed by other researchers (Nir *et al.*, 2007). High versus low firing rate neurons have been found to show other physiologic differences in recent years (Mizuseki *et al.*, 2009; Buzsáki & Mizuseki, 2014; Grosmark & Buzsaki, 2016; Watson *et al.*, 2016) and so further information may be extracted using this spike-power correlation difference.

As brain state varies, the spike-LFP correlation also changes. In particular nonREM sleep has the strongest cortical LFP oscillations, in the form of delta waves (or “slow waves”) and in that state the mid-frequencies spanning from 5-50Hz are all positively correlated with spike rates, unlike during WAKE or REM. Delta power remains negatively correlated with spike rates in all states and with all bin sizes, while 50-180Hz power remains positively correlated in all states. While the full population correlation remains positive in this band, we showed that single neurons could switch their preference for spiking with the 50-180Hz band across brain states. Spiking activity of putative pyramidal cells typically co-varied with gamma power, but a minority showed an inverse relationship. In most cases, the negative correlation was observed during nonREM sleep. In future studies, it will be interesting to determine functional differences between neurons that do or do not switch preference for gamma band.

In contrast to the gamma band, delta, sigma and beta band power were anti-correlated with spike rates during the wake state. Theta frequency was the only low frequency band which displayed a positive correlation with spike rates. Theta LFP may or may not be generated in the neocortex and the positive coupling likely corresponds to entrainment of neocortical neurons by the hippocampal-entorhinal projections (Sirota *et al.*, 2008).

We used two additional complementary methods to assess the relationship between spiking and LFP: LFP phase modulation and spike-triggered wavelet spectra. In wake and REM sleep, the high gamma band in the 50-180Hz range showed preferential modulation of spike timing, while in nonREM sleep, positive spike-LFP coupling expanded down to the delta frequency band. Furthermore, epochs of higher power had stronger phase modulatory effects than lower power epochs for most frequency bands. The spike triggered wavelet-averaging approach showed similar power distributions to bin-based methods and peaks were also shared with phase modulation metrics, lending each credence by dint of their convergence.

### Excitatory inhibitory coordination

The spike triggered LFP average analysis also showed a consistent effect of excitatory-inhibitory cell firing patterns where E cells tend to precede I cells by 1-3 milliseconds. This is a novel finding but which is consistent with both recent intracellular findings on post synaptic potential relative timing (Haider *et al.*, 2016) and with models of gamma oscillations (Börgers *et al.*, 2008; Isaacson & Scanziani, 2011), This regularly timed excitatory-inhibitory timing may also relate to our consistent finding of increased pI to pE activity ratios with higher broadband gamma powers. Much evidence indicates the role of parvalbumin-positive interneurons in gamma oscillations (Cardin *et al.*, 2009; Buzsáki & Wang, 2012) and optogenetic studies suggest that the pI neurons we record are largely parvalbumin positive (Stark *et al.*, 2013). The change in excitatory-inhibitory balance with higher gamma band power observed here may be due to employment of an inhibition-related mechanism to generate these oscillations, and that mechanism maybe PING-like (Börgers & Kopell, 2003) (Stark *et al.*, 2013), (Moca *et al.*, 2011; Buzsáki & Wang, 2012). Thus, the 50-180Hz band may represent not just increased spiking overall, but actually may signify increased engagement of inhibitory interneurons and therefore a fundamentally different local network state.

### EMG contamination of LFP

An often neglected by important but artefactual source of high gamma power is electromyographic (EMG) artifacts, volume-conducted from head and neck muscles (Whitham *et al.*, 2007). EMG artifacts are ubiquitous in all animals, including humans, yet spectral or wavelet measures alone cannot distinguish them from neuron-induced transmembrane currents. We found a positive correlation between EMG magnitude and LFP power down to 50 Hz frequency. Since running speed and the power of gamma oscillations may co-vary (Bragin *et al.*, 1995; Ahmed & Mehta, 2012), it is not clear to what extent the increased power reflects true LFP oscillations versus EMG contamination. The large difference in power at >200 Hz between REM sleep, with it’s generalized muscle atonia (and occasional EMG twitches) and waking suggest that a large part of the high frequency power is due to EMG. Importantly, the large waking LFP power above 200 Hz was not reflected in the spike-LFP coupling measure, in support of non-neuronal nature of this band. We recognized and quantified EMG from the zero-time high coherence across all recording electrodes (Watson *et al.*, 2016). It is important to note that zero-phase lag synchronized high frequency gamma activity (Vicente *et al.*, 2008) recorded across structures has been suggested to reflect various cognitive events, such as memory recall and decision-making (Stujenske *et al.*, 2014; Yamamoto *et al.*, 2014). In future studies, multiple electrodes should be used to separate possible physiological coupling of gamma oscillations from EMG-meditated contamination.

### Maximizing interpretability of LFP

Given the spike waveform and EMG contaminants of LFP signals, moving into the future further work should be done to make maximally-interpretable LFP signals. A combination of technical and analytical approaches can help in rodents or other systems with greater control in order to assist the field in ways that allow improved interpretation of both rodent and human data. First, simultaneous recording of LFP and spiking of isolated pyramidal cells and interneurons confers a greater ability to assess both the involvement of different cell populations in the coordinated activity reflected in the LFP and the directionality of communication. Second prior knowledge of anatomical connectivity may assist in placing the recording electrodes in appropriate layers for testing hypotheses about how excitation propagates through the network. Next, prevention or removal, or at least consideration, of spike and EMG contamination of high-frequency LFP power and phase is an absolute necessity. Effective removal of these contaminants by appropriate offline analysis can be of great benefit.

Assuming that the above conditions are met, sophisticated multivariate methods are available for teasing out cause-effect relationships, including directed coherence, Granger causality and dynamic causal modeling (Pereda *et al.*, 2005; Friston *et al.*, 2013), and independent component analysis (Fernandez-Ruiz *et al.*, 2012) might be harnessed to extract important functional physiological interactions between different parts of neural circuits. Many of these techniques are deployed with the intention of overcoming limitations in the measurements, but it is important to bear in mind that these computational methods are only as good as the physiological recordings. No amount of mathematical intervention can rescue insufficient input data. On the other hand, these promising methods may be further refined and placed on firmer footing if they can be tested on data sets with less physiological ambiguity.

### Conclusions

We have illustrated here variance in the spike-LFP coupling across cell types, states and frequency bands. We find support for a possible excitatory-inhibitory joint mechanism for gamma band oscillations. We also suggest metrics to better predict spiking activity from LFP signatures. Yet, much clearly remains to be understood. We believe that the best route forward may therefore be to advance our general knowledge of LFP relationships with spiking by deploying more advanced recording techniques improved analytical methods and experimental perturbation of specific circuit elements to disentangle the activities of distinct neural populations, their emergent interactions in the gamma frequency band, and the relationships between their dynamics with behavior.

## Acknowledgments

This work was supported by National Institutes of Health Grants (MH54671; MH102840; MH107662), the Mather’s Foundation, The Human Frontier Science Program and the National Science Foundation (Temporal Dynamics of Learning Center Grant SBE 0542013), The Leon Levy Foundation and the Brain Behavior Research Foundation. We thank Norman Itzik, Antonio Fernández-Ruiz, Kenji Mizuseki, Antal Berényi, and members of the Buzsáki laboratory for discussions and feedback on the manuscript. We especially thank Dr. Jennifer Gelinas and John Palmer Greene for help with collection of original recorded data.

## AUTHOR CONTRIBUTIONS

BOW and GB deigned experiments. BOW conducted experiments. BOW and MD performed analyses. BOW and GB composed the manuscript

## ABBREVIATION LIST

LFP: Local field potential

EMG: Electromyographic signal

MUA: Multi-unit activity

REM: Rapid Eye Movement sleep

pE: Putative excitatory unit

pI: Putative inhibitory unit

## Supplementary Figures

**Supplementary Figure 1.**
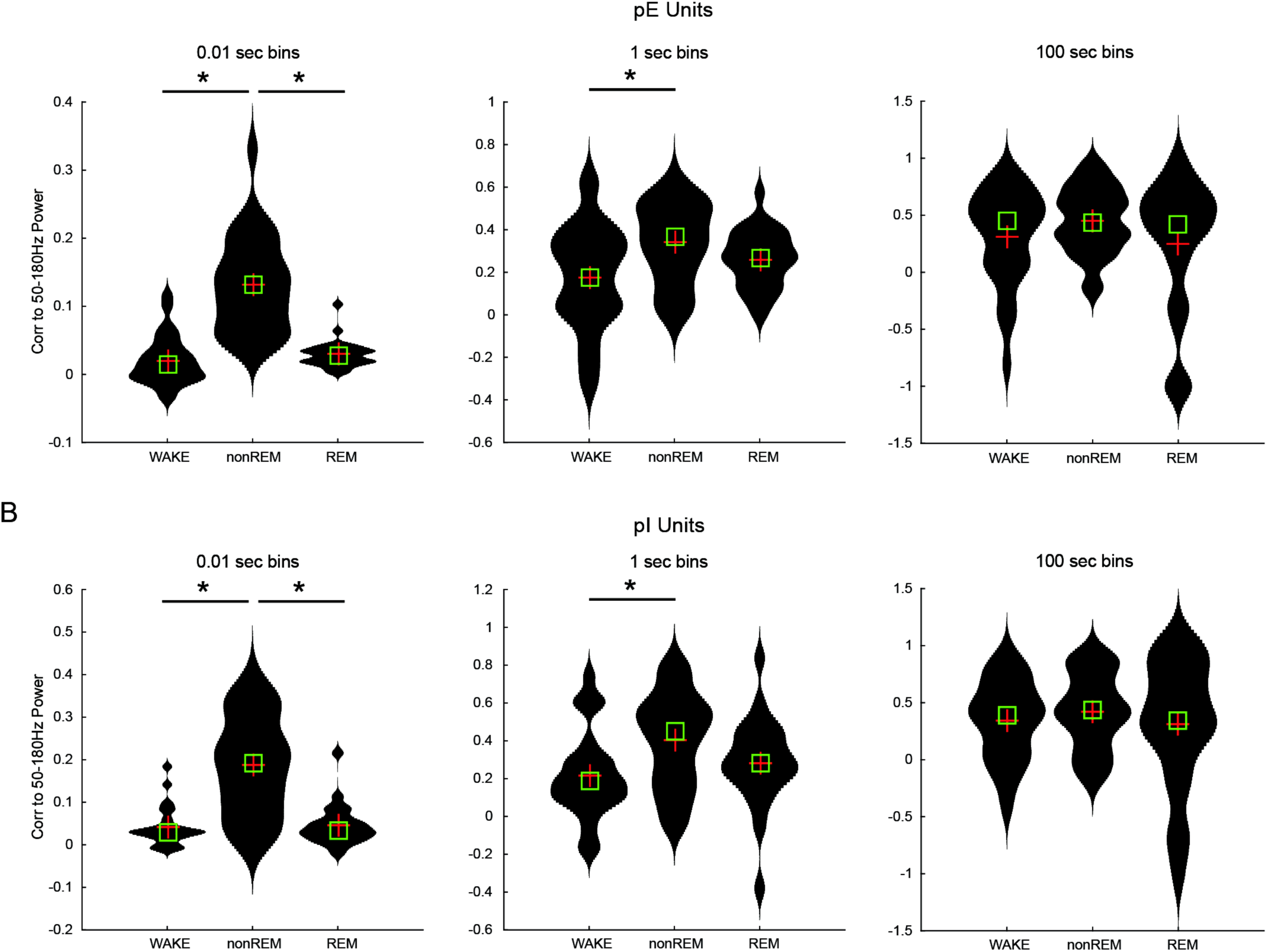
Variance by state and time bin in the correlation of population spike rate versus broadband gamma. A. Putative excitatory unit cumulative population spiking rate (sum of spikes across all simultaneously recorded units) versus 50-180Hz broadband gamma power on a per-bin basis. Bin sizes are 0.01s, 1s and 100s in left, center and right panels respectively. Within each panel, each column shows the distribution of session-wise Pearson r’s for WAKE, nonREM and REM. At 0.01s and 1s bins, the correlation of spike rate to 50-180Hz power is significantly greater for nonREM than it is for WAKE and REM (difference with p<0.05, indicated by asterisk). With 1 second bins the nonREM rate-broadband gamma correlation is higher than WAKE and with 100 second bins there are no significant differences in correlations between states. B. Same as A but for putative inhibitory units. State-wise significant difference profiles by bin size are not different between pE and pI units.

**Supplementary Figure 2.**
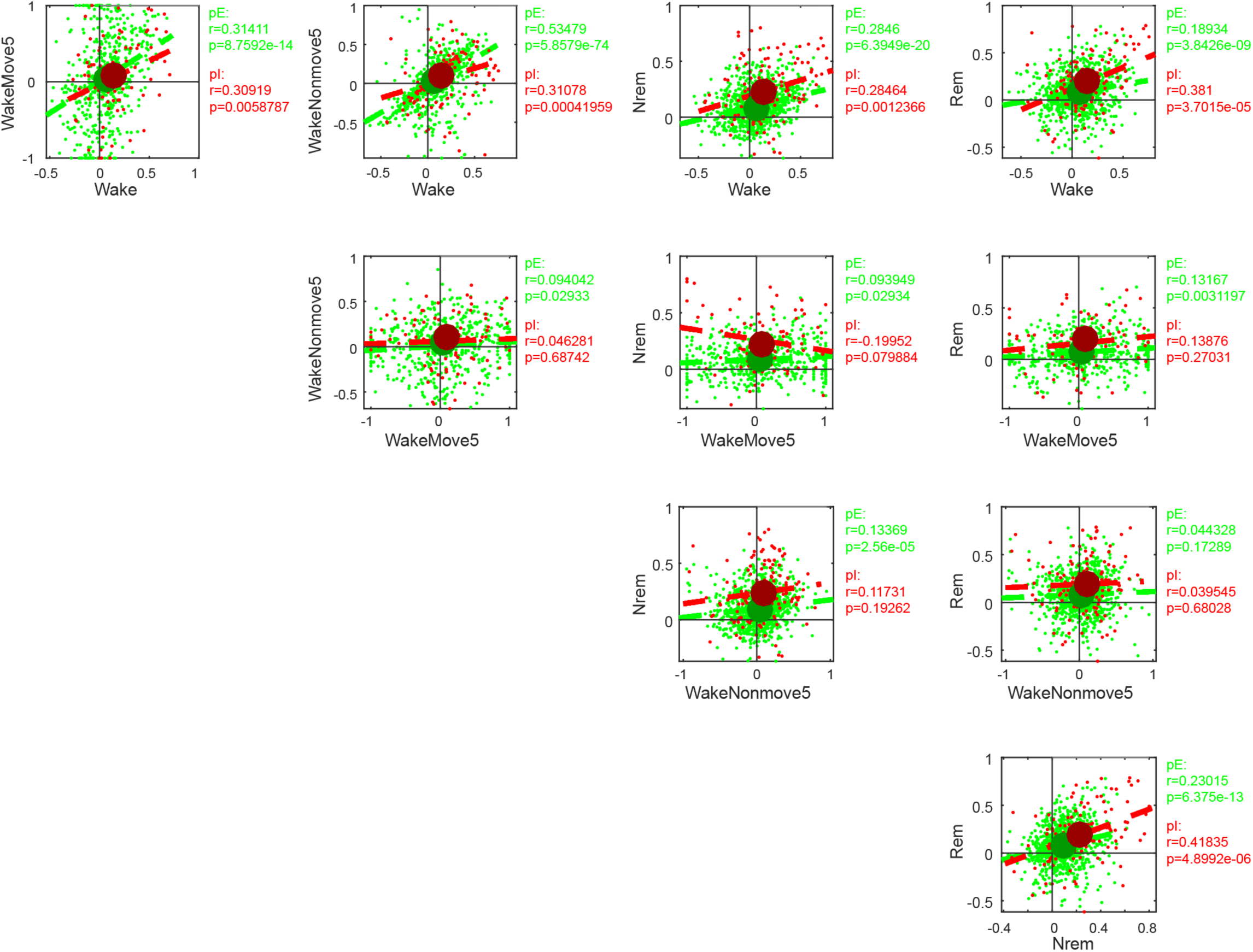
Variance of spike rate-broadband gamma coupling within each unit depending on state. Single neurons (green or red dots) are shown in plots illustrating Pearson correlations of that neuron’s spike train versus broadband high gamma power for pairs of states (ordinate and abscissa). Large green and red dots show population averages for pE and pI units respectively. Note that while the population averages show a positive correlation for R values in one state versus R values in other states, there is a large variance around this trend with units frequently switching their modulation by broadband high gamma according to state.

**Supplementary Figure 3.**
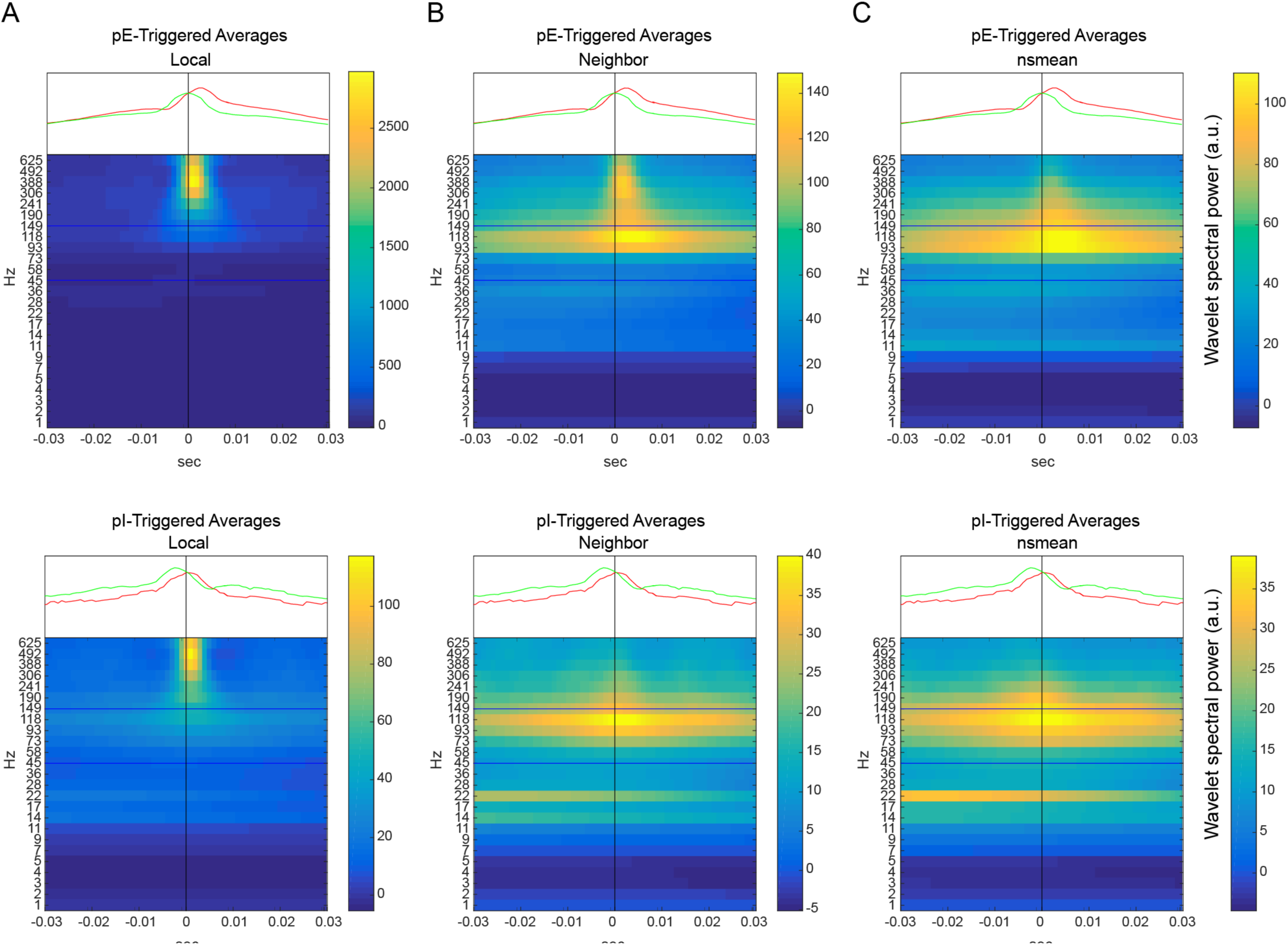
Spike-triggered wavelet spectra and pE and pI unit spiking broken down by cell type (pE versus pI) and proximity of LFP channel to unit channel. A. pE (top) and pI (bottom) unit-triggered spectra on the channel local to the unit spike itself. Note that for both pE and pI units there is a substantial increase in spectral power above 300Hz. B. pE and pI unit-triggered spectra for LFP from the neighbor shank 200um away from the shank on which the unit is recorded: for pE units some degree of 300Hz power increase remains, while it does not for pI neurons. Note that the high gamma broadband range is also increased in power around spiking. C. When wavelet spectra from yet more distant channels are included (non-shank mean of LFP of 1 site per non-neighbor shank (nsmean)), the 300+Hz power disappears and only gamma-band power remains elevated. In each plot the spike rates of pE and pI units outside of the trigger unit are also shown in green and red line-plots. Note that pE units tend to fire approximately 3ms prior to pI units across all states and across pE or pI triggering states.

**Supplementary Figure 4.**
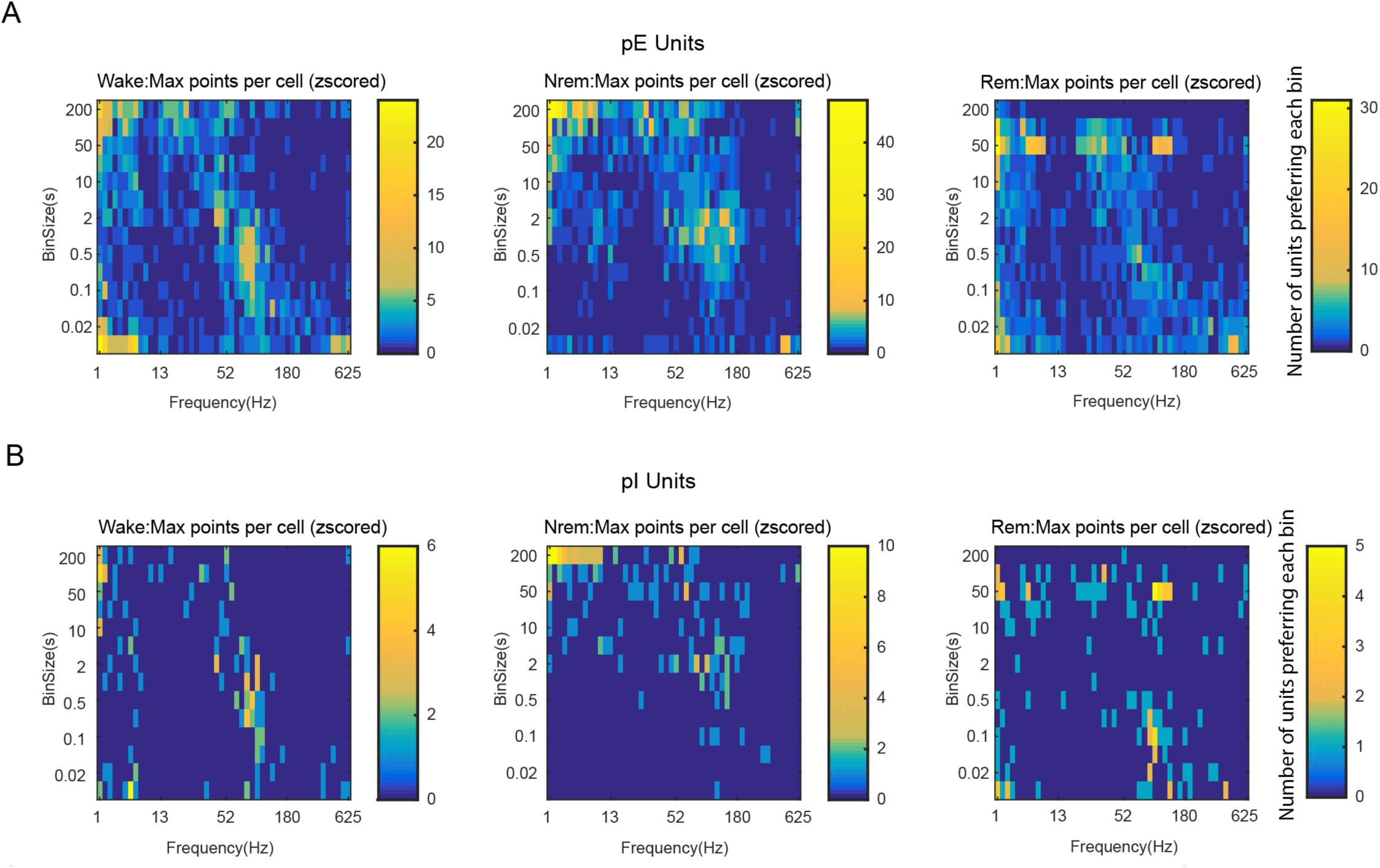
A. Two-dimensional histograms broken down by brain state of maximal Pearson r values across the bin sizes and frequency bands for each pE unit in the dataset. The 50-180Hz band again dominates with most cells preferring bins in this band, but many neurons also correlate with lower frequency power ranges as well. Overall maximal density of unit preference is for the combination of 0.5 second bins and 90Hz LFP. B. Same analysis as panel A but for pI units

**Supplementary Figure 5.**
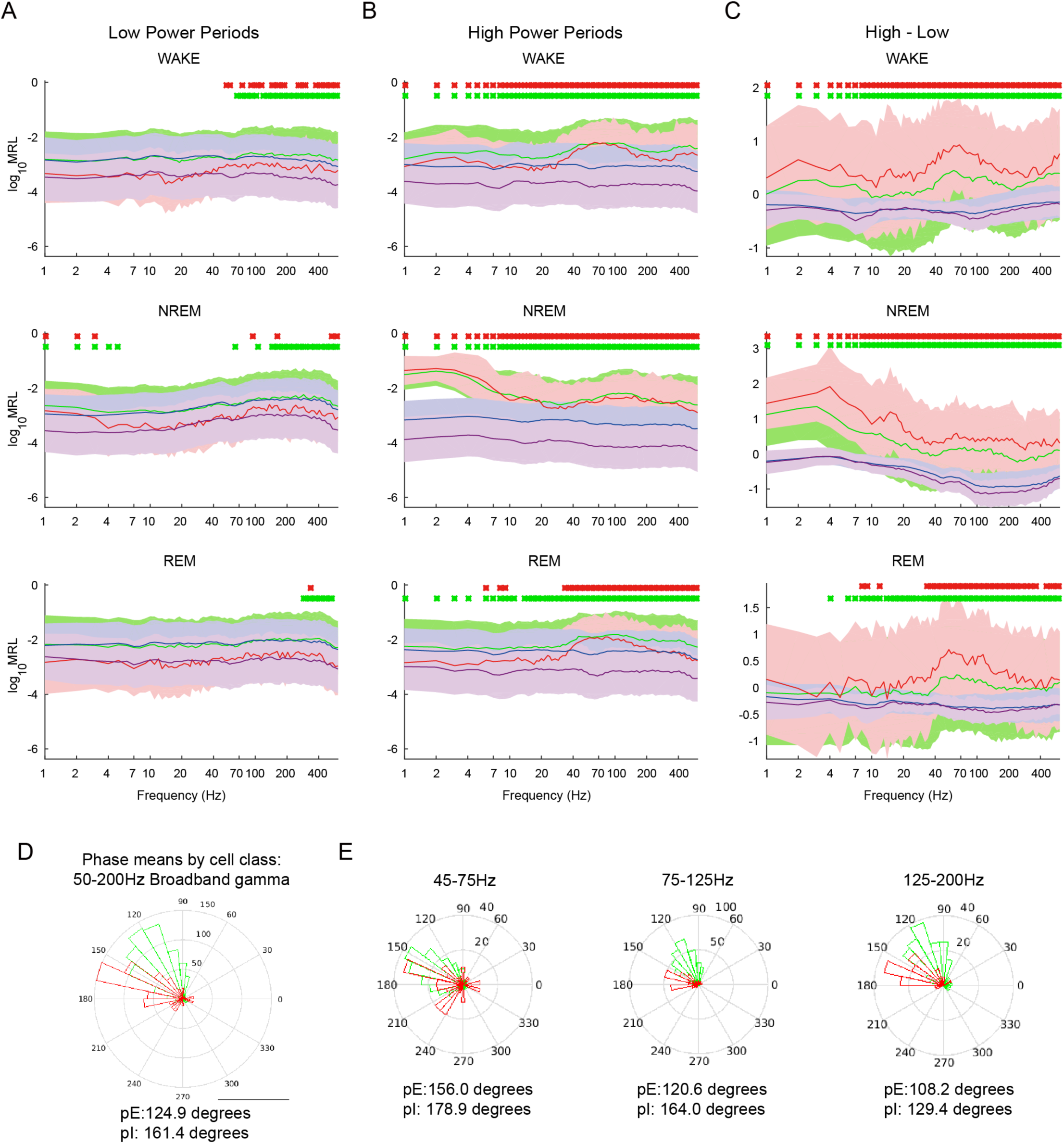
Phase modulation increased with higher amplitude oscillations than with lower frequency band oscillations, and pE and pI units show differential phase preference to broadband gamma. A. Log of mean resultant vector lengths averaged across all pE units and pI units in the dataset, across all frequencies, stratified by state, only for low power times in each oscillation, defined as less than zscore -0.5 for at least 3 cycles of oscillation time. pE units shown in green, pE shuffles in blue, pI in red and shuffled pI in purple. Green and red dots above specific bands indicate bands of significant difference based on Wilcoxon signed rank test - green dots for pE units, red dots for pI units. B. Same as A, but for high power times only, defined as continuous epochs of power greater than z- score 0.5 for at least 3 cycles of oscillation time. Note greater significant phase modulation as indicated by dots above plots C. Difference between high and low power modulation curves for each frequency band. Note that, higher power periods preferentially phase modulate spiking more than low power periods, consistent with a causal relationship between spike timing and LFP phase, especially for broadband gamma but also for delta band in nonREM sleep. D. Mean phase differences for pE (green) versus pI units (red) in the broadband gamma range. Shown are circular histograms of average per-unit mean phase. E. Same as D but for narrower frequency bands, indicating a consistent 20-25 degree lag of I units behind E units – or about 0.4-1.5ms over the various frequencies.

**Supplementary Figure 6.**
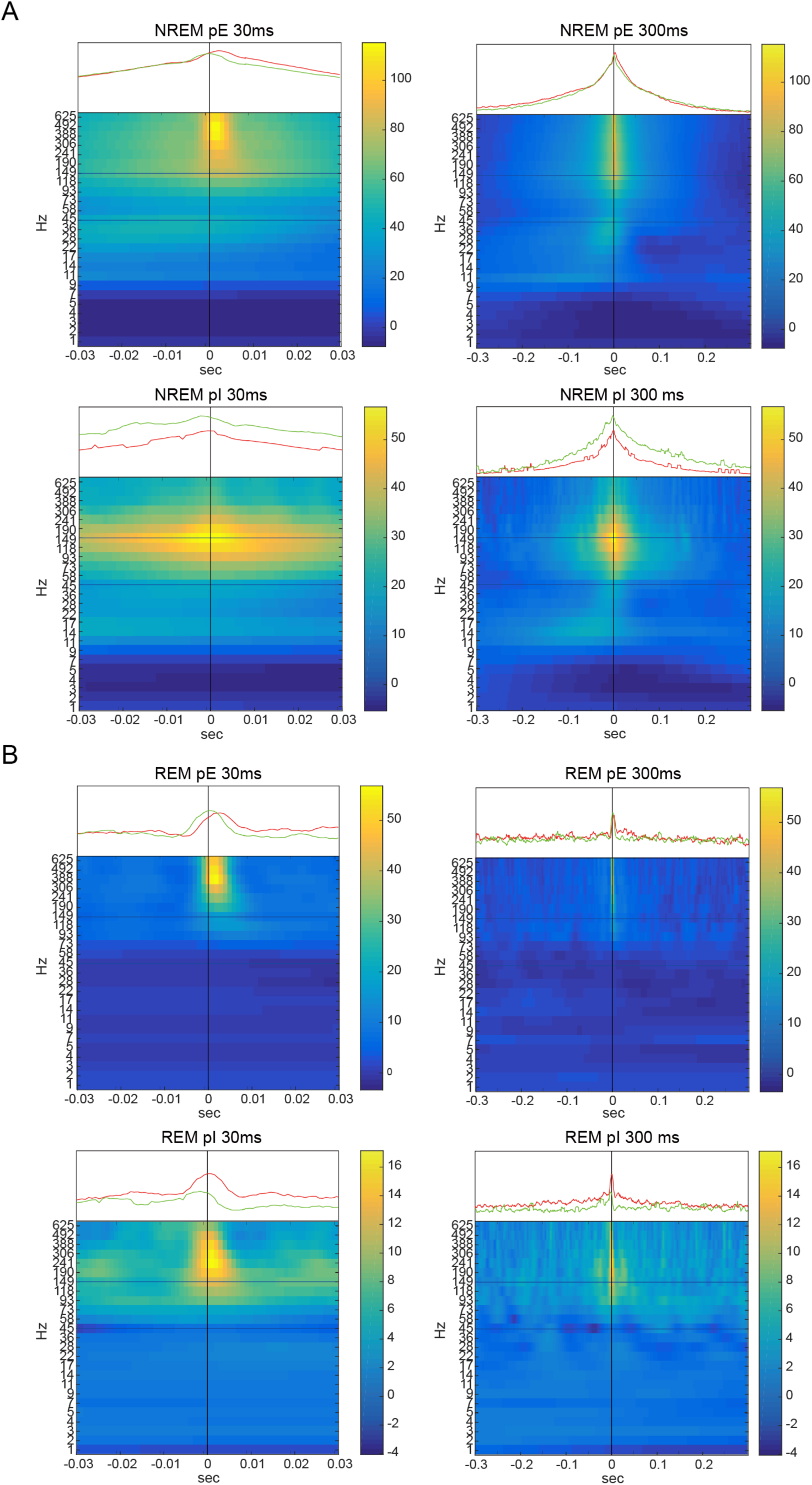
Spike-triggered wavelet spectra from neighboring shanks for pE and pI units in the non-WAKE states of nonREM and REM sleep. Green and red lines above each plot show spike-triggered firing of the surrounding pE and pI populations near each trigger unit (excluding units recorded on the same shank).

**Supplementary Figure 7.**
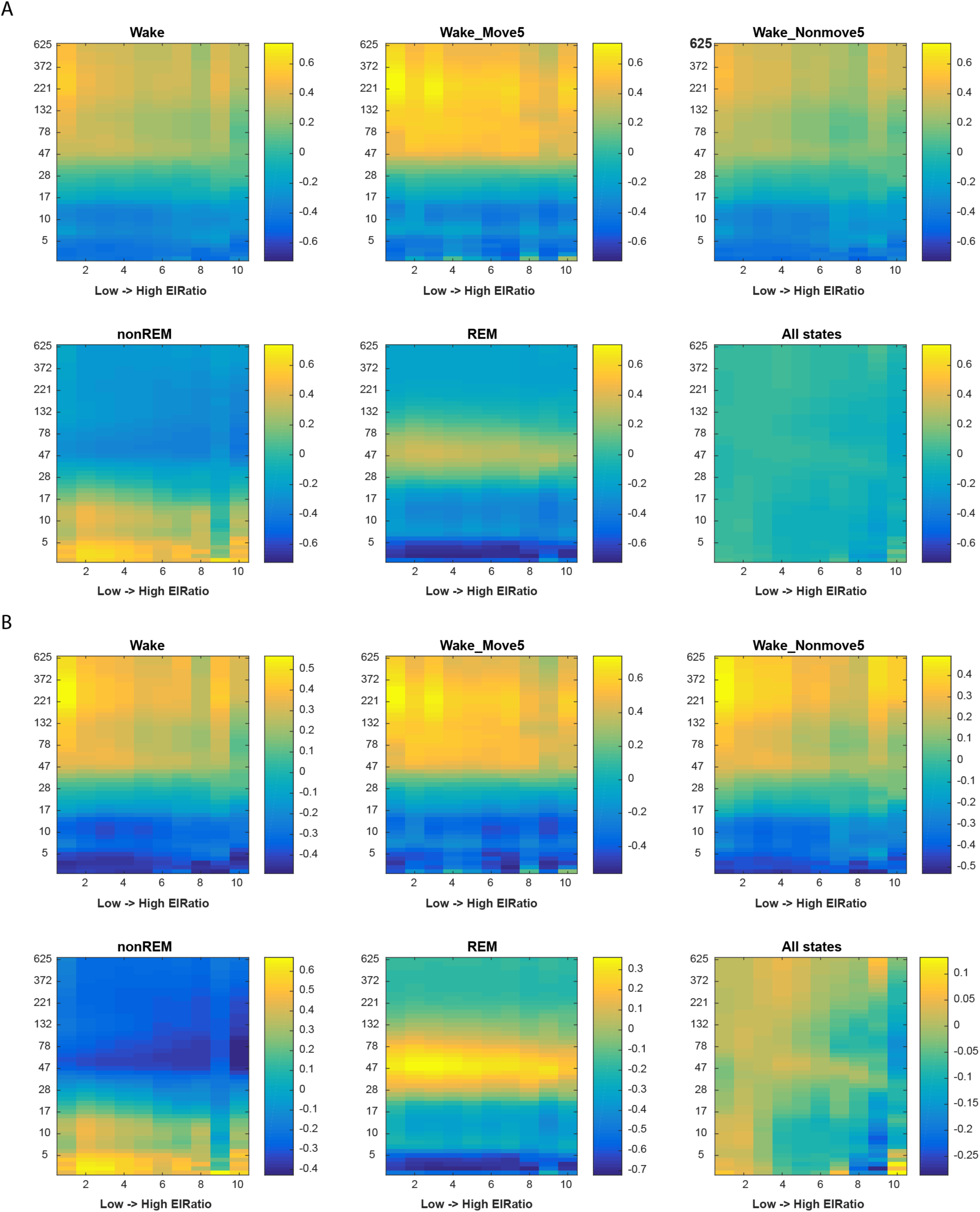
LFP correlates of EI Ratio: LFP spectral powers per second-long bin after those bins were ranked into high to low deciles of pE to pI unit relative activity. A. Raw powers of spectra averaged over the entire population, on a common power-color lookup table across all states in order to allow for relative modulation of LFP by EI Ratio by state. B. Individualized power-color look-up tables for each state in order to reveal within-state relative modulation of LFP spectra by EI ratio, prior to z-scoring as in Figure 7. Note muting of modulation when including All states, rather within-state modulation appears to be a clear feature which can be obscured with intervening state transitions.

**Supplementary Figure 8.**
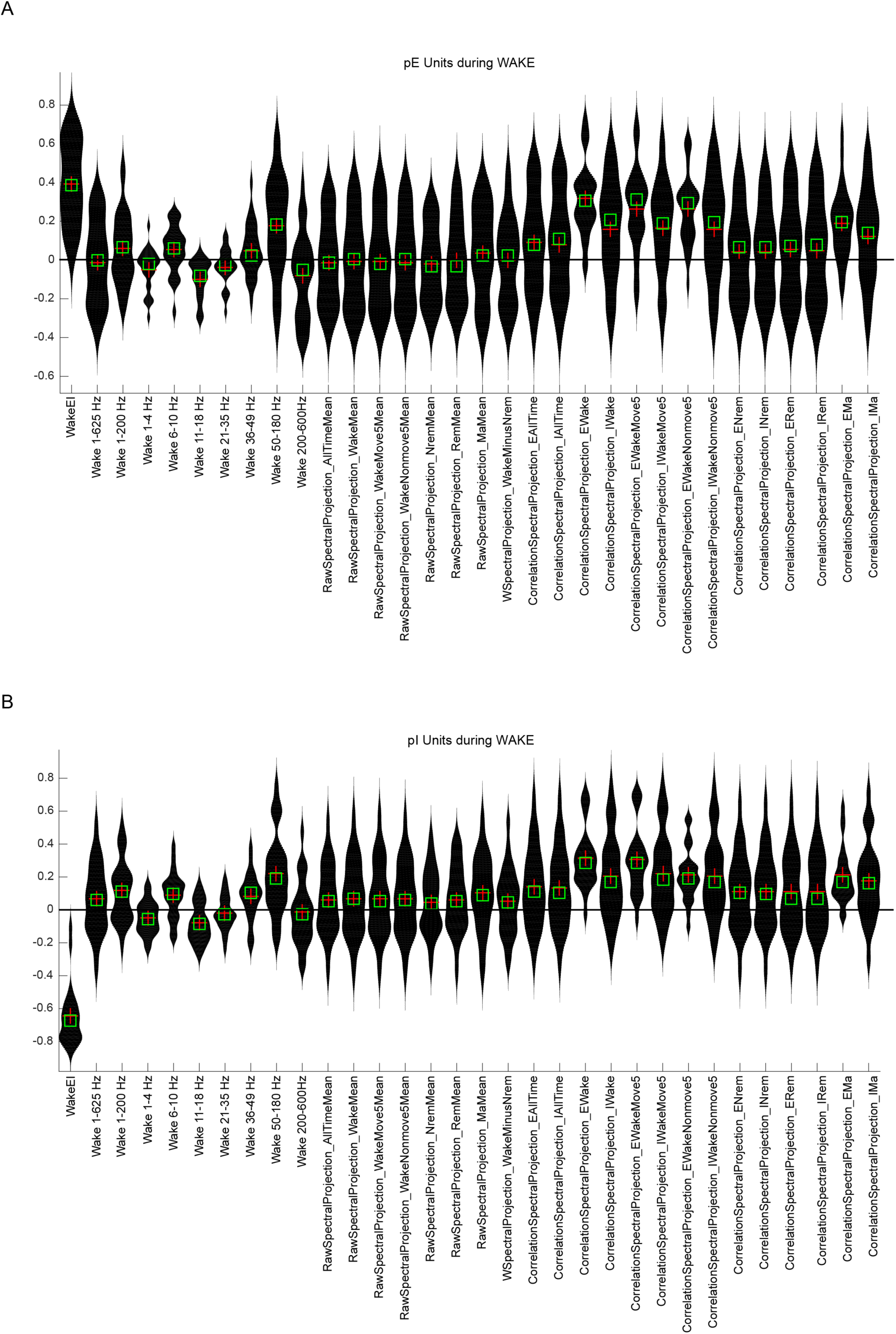
Correlates of pE and pI unit population spiking. A. 31 tested predictors of spike rates, all but the first, EIRatio, are LFP-based. All correlation measured carried out during the WAKE state to predict spiking in that state. EI ratio is on the far left. The next 9 measures are spectral band powers. The metrics labeled “RawSpectralProjection_…” represent dot product projections of averaged state-wise spectra (as in Figure 2B) onto the measured spectrum of each time bin. There is next a special interest case of the wake state spectrum minus the nonREM state spectrum. Finally, are the correlation spectral projections, gathered from correlations of pE or pI unit types in various states, as labeled. The correlation spectrum for pE units in WAKE or WAKE-related states predicted firing rates most strongly and with the least negative correlation compared to other predictors.

